# Thermal oscillations enable reshuffling of genetic material in a primitive cell cycle

**DOI:** 10.1101/2021.04.01.438038

**Authors:** Roger Rubio-Sánchez, Derek O’Flaherty, Anna Wang, Francesca Coscia, Lorenzo Di Michele, Pietro Cicuta, Claudia Bonfio

## Abstract

Self-assembling single-chain amphiphiles available in the prebiotic environment likely played a fundamental role in the advent of primitive cell cycles. However, the instability of prebiotic fatty acid-based membranes to temperature and pH seems to suggest that primitive cells could only host prebiotically-relevant processes in a narrow range of non-fluctuating environmental conditions. Here we propose a novel primitive cell cycle driven by environmental fluctuations, which enable the generation of daughter protocells with reshuffled content. A reversible membrane-to-oil phase transition accounts for the dissolution of fatty acid-based vesicles at high temperatures, and the concomitant release of genetic content. At low temperatures, fatty acid bilayers reassemble and encapsulate reshuffled genetic material in a new cohort of protocells. Notably, we find that our disassembly/reassembly cycle drives the emergence of functional RNA-containing primitive cells from parent non-functional compartments. Thus, by exploiting the intrinsic instability of prebiotic fatty acid vesicles, our results point at an environmentally-driven tunable primitive cell cycle, which supports the release and reshuffle of protocellular genetic and membrane components, potentially leading to a new generation of protocells with superior traits. In the absence of protocellular transport machinery, the environmentally-driven disassembly/assembly cycle proposed herein would have supported genetic content reshuffling transmitted to primitive cell progeny, hinting at a potential mechanism important to initiate Darwinian evolution of early lifeforms.

## Introduction

The emergence of primitive cell cycles represents a step towards the generation of model protocells. Inspired by modern biology, biochemists and synthetic biologists aiming to mimic evolution have long tried to understand how minimal cells could recursively undergo *model* cell cycles of growth and division,^1,2^ while continuing to sustain compartmentalised biochemical processes. One unresolved difficulty with proposing *primitive* cell cycles is that prebiotic components should be employed, and the identified model conditions should be compatible with the prebiotic environment and processes.

Among the prebiotically plausible biological building blocks, fatty acids were likely major components of primitive membranes.^3–6^ Recent reports have demonstrated how fatty acid-based compartments can host prebiotically-relevant reactions, including activation of building blocks^7^, metal-based catalysis^8^ and nonenzymatic RNA replication.^4,5,9^ However, the limited stability of fatty acid-based protocells to changes in temperature, pH and ionic strength^10–12^ needs to be taken into account when one aims at expanding the repertoire of compartmentalised prebiotic processes and identifying plausible pathways of primitive cell replication.

Fatty acid-based protocells have been shown to undergo cycles of growth and division.^13–15^ Specifically, the growth of large multilamellar vesicles can be achieved by addition of fatty acid micelles, resulting in long thread-like bilayer structures that, upon shearing, divide into smaller vesicles with no significant loss of protocellular content.^15^ Such a model of primitive cell cycle relies on two assumptions: i) that the required environmental conditions are not subject to fluctuations, as not to destabilise fatty acid vesicles, and ii) that content reshuffling, product dilution or substrate uptake is not required for encapsulated prebiotic reactions to work. However, natural temperature and pH gradients would have easily formed both in sunlit aqueous surfaces and hydrothermal environments on early Earth^19–21^ and, likely, would have been required to support most prebiotic (bio)chemical processes, including the synthesis^16^ and polymerisation^17,18^ of life’s building blocks. Also, in the absence of modern trafficking biosystems to rely upon, accumulation of products or lack of substrates could have inhibited compartmentalised protocellular processes.^22,23^

In this study, we unravel a new, prebiotically-relevant model cell cycle, in which thermal oscillations drive the formation of new cohorts of fatty acid-based protocells and the reshuffling of their encapsulated material. At high temperatures, unilamellar vesicles collapse at first into multi-layered structures, then into lipid droplets, concomitantly releasing their protocellular content. At low temperatures, the self-assembly of fatty acid protocells *via* membrane budding from the surface of lipid droplets yields daughter protocells with re-encapsulated protocellular material. Such lipid phase transitions, triggered by thermally-induced pH fluctuations, can be modulated by the presence of relevant building blocks, like nucleotides and peptides. Thus, by exploiting the thermal instability of prebiotically-relevant lipid vesicles, our work proposes a novel geochemically plausible primitive cell cycle, underpinned by the reversible pH-driven assembly and disassembly of the protocellular state, and addresses for the first time the fundamental role of protocellular (genetic) content mixing and reshuffling.

## Results

### Reversible and tunable lipid phase transitions

Moderate thermal gradients, such as those naturally generated in hydrothermal environments or sunlit shallow ponds, were shown to support the amplification of functional nucleic acid strands,^24,25^ as well as the accumulation and self-assembly of prebiotic amphiphiles (between 5°C and 50°C).^26^ However, previous studies on the permeability of primitive cells showed that loaded vesicles, made of short-to-medium chain fatty acids, do not retain oligonucleotides when exposed to high temperatures (above 50°C).^12^ While heat-stable lipid mixtures were identified, the reported thermal instability of certain prebiotic vesicles begs the following question: What mechanism could cause the release of the encapsulated content at high temperatures?

To shed light on the behaviour of fatty acid vesicles upon heat exposure, we sought to explore whether alterations in membrane morphology might occur and be responsible for the leakage of the encapsulated content at high temperatures. A buffered solution containing extruded myristoleic acid vesicles was heated up to 95°C and changes in membrane fluidity and turbidity were detected by fluorescence and UV-vis spectroscopy. Myristoleic acid was chosen as a proxy for self-assembling amphiphiles^9^. The lipophilic probe Laurdan was embedded in the lipid bilayer to provide information on changes in membrane structure and dynamics.^13^ Notably, at high temperatures we observed a sharp increase in turbidity (420 nm), concurrent with a drastic decrease in the fluorescence intensity ratio of Laurdan emission maxima (496/426 nm) (Figure 1a and S1). Remarkably, vesicles prepared with prebiotically-relevant amphiphile mixtures (decanoic acid and decanol, 2:1 ratio)^3^ showed similar absorbance and fluorescence profiles (Figure S2). Such results point towards a major heat-driven alteration in lipid packing.

**Figure 1.**
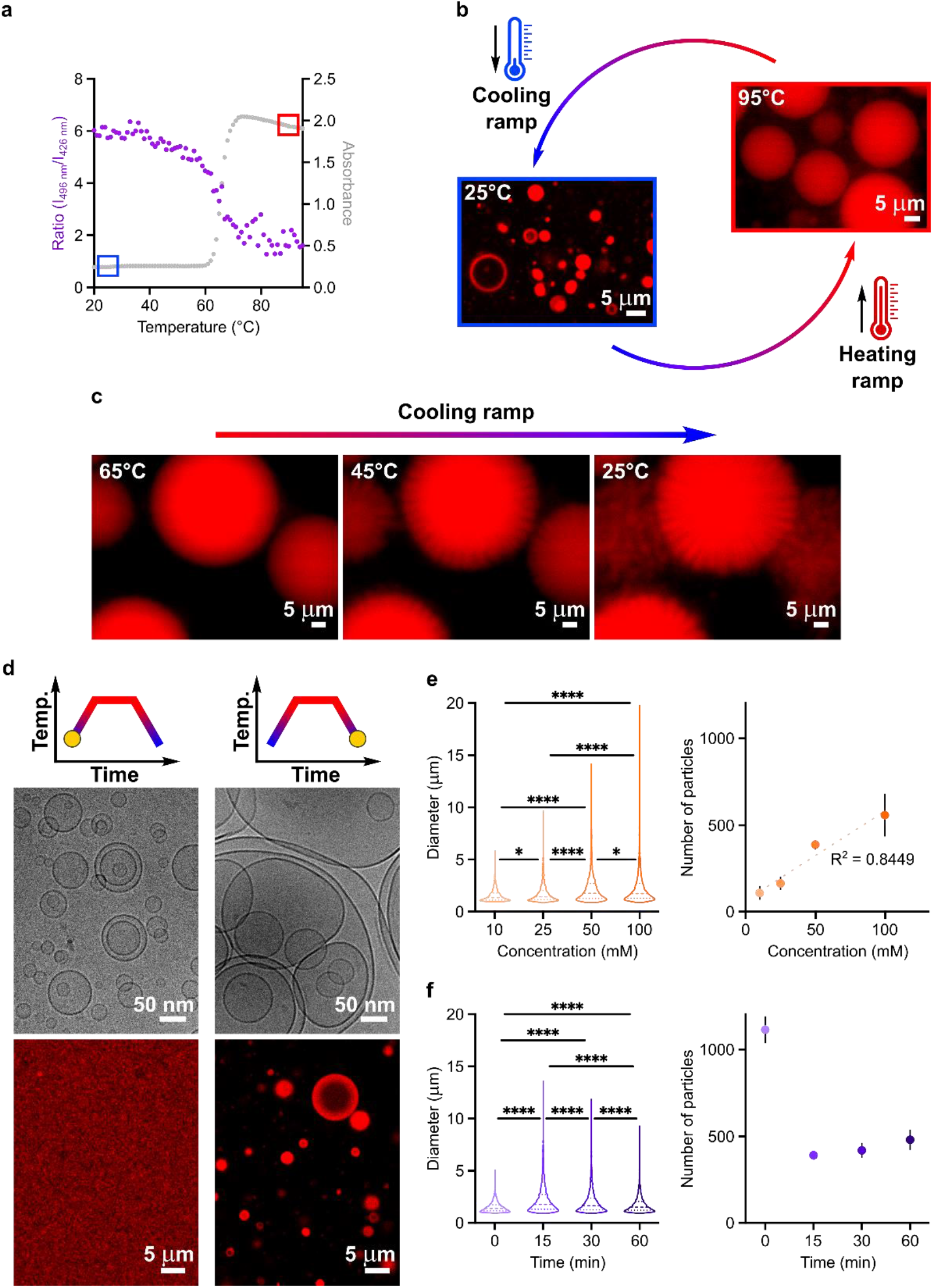
Thermal cycling drives reversible and tunable fatty acid phase transitions. a) Overlapped profiles of turbidity and fluorescence intensity as a function of temperature for 100 nm-radius vesicles made of 50 mM myristoleic acid (in 200 mM Tris-HCl, pH 7). Absorbance (grey) is monitored at 420 nm, whereas Laurdan fluorescence (purple) is monitored at λ_exc_ = 426 nm and 496 nm. b) Schematic representation of thermal cycles explored in this study. Microscopy images after the heating ramp (95°C, hot-stage epifluorescence microscopy images) and after the cooling ramp (25°C, confocal microscopy images) are shown for myristoleic acid vesicles. c) Membrane budding observed during the cooling ramp on the surface of myristoleic acid droplets by hot-stage epifluorescence microscopy. d) Cryogenic electron microscopy (top) and confocal microscopy (bottom) images collected before and after thermal cycles for myristoleic acid vesicles. e) Analyses of myristoleic acid samples showing how the size (left) and number (right) of the next-generation population of vesicles are affected by lipid concentration. e) Analyses of myristoleic acid samples showing how the size (left) and number (right) of regenerated vesicles are affected by heating time. Analyses in d) and e) were performed on collected confocal microscopy images for 100 nm-radius vesicles made of 50 mM myristoleic acid vesicles in 200 mM Tris-HCl, pH 8 after thermal cycling. Upon cooling the samples down to 25°C, vesicles were allowed to re-equilibrate for 1 h. Statistical significance was assessed using the one-way ANOVA test, n = 10 replicates. Statistical values obtained for: * P = <0.05, **** P = <0.0001. n = 3 for data in a) and d). Center dashed line represents median; dotted lines represent upper and lower quartiles.

To identify the cause of increased turbidity, we modelled^27^ the light-scattering profile that would be expected for extruded unilamellar vesicles upon heat exposure. The calculated optical density value is in perfect agreement with our experimental observations for decanoic acid-based vesicles (Figure S3). Even though a small increase in optical density could either be accounted for by an increase in vesicle radius or lamellarity (Figure S3a-b), the observed increase of absorbance above 0.5 can only be attributed to the formation of larger dense structures with high internal lipid fraction (Figure S3c). Our model thus suggests that heat induces not only an increase in vesicle size, but also in lipid density. In order to elucidate the effect of heating on the morphology of fatty acid membranes, we turned to hot-stage epifluorescence microscopy. Along the heating ramp (25°C→95°C, at a rate of 0.1°C·s^−1^), myristoleic acid vesicles exhibit an increasingly dynamic behaviour (Movie S1). Unexpectedly, the surface area of lipid vesicles increases with temperature, resulting in elongated structures, which eventually collapse into micron-sized lipid droplets at high temperatures (Figure 1b and S4). Kept at 95°C for 15 minutes, fatty acid droplets coalesce into larger structures. Then, upon cooling down to 25°C, membrane budding from the water-oil interface results in the generation of daughter fatty acid vesicles (Figure 1c). These findings are consistent with both our modelling and experimental spectroscopic data.

As illustrated in their phase diagrams, fatty acids form membranes spontaneously in aqueous solutions, in a narrow pH window around their apparent pK_a_.^28^ At high pH, fatty acids are fully deprotonated and form micelles, whereas, at low pH, they are fully protonated and form an oil phase.^28^ Additional lamellar structures, as well as cubic and hexagonal phases, are observed at intermediate pH values or higher lipid content.^29,30^ As the pK_a_ of most functional groups vary appreciably with temperature,^31^ the reversible phase transition observed for myristoleic acid vesicles likely results from pH fluctuations during heating and cooling ramps. In further support to our hypothesis, the temperature-dependence of pH for a number of biological buffers was evaluated by means of UV-vis spectroscopy in the presence of fluorescein, a pH-sensitive probe (Figure S5). While the pK_a_ of phosphate was only mildly affected by heat, the pH of solutions containing Tris-HCl dropped by two units when heated from 25°C to 95°C. Such results suggest that myristoleic acid vesicles prepared in Tris-HCl buffer are destabilised upon heating, *i.e.,* when the pH becomes too acidic (below 6.5), and convert to lipid droplets. At high temperatures, the local increase in interfacial tension, due to the high concentration of protonated fatty acids, likely leads to the high-energy lipid surfaces coalescing into larger droplets. Upon cooling, as deprotonation of fatty acids in the outermost leaflet of lipid droplets could lead to increasing curvature, undulating folds occur on droplet surface, resulting in membrane budding. As such, a series of experiments was performed to interrogate lipid dynamics and phase transitions systematically for both myristoleic acid and decanoic acid-based membranes under different conditions. The reversible fatty acid vesicle-to-lipid droplet conversion occurs in most biological buffers, and in combinations thereof, as well as in self-buffering conditions, thus indicating that such lipid phase transitions could have naturally occurred in a wide range of primordial buffered and unbuffered environments (Figure S6-8). To explore the tunability of phase-transition temperatures, variations of initial pH and ionic strength values, buffer and fatty acid concentrations were also tested (Figure S9-13). Intriguingly, the addition of peptides and nucleotides, as prebiotically-relevant building blocks, shifts the transition temperature to lower values in a charge-dependent manner, likely due to ionic interactions with lipid headgroups affecting the bilayer’s hydrogen bonding network (Figure S10). These results indicate that temperature gradients generate pH fluctuations in most buffered and unbuffered environments and drive reversible fatty acid phase transitions, which can be altered by the presence of life’s biological building blocks.

The spontaneous self-assembly of fatty acid membranes generally results in polydisperse multilamellar structures and, from those, unilamellar vesicles are then formed by extrusion.^28^ Our confocal and cryogenic electron microscopy data reveal that, when 50 mM extruded small unilamellar myristoleic acid vesicles undergo thermal cycling (25°C→95°C→25°C, at a rate of 0.1°C·s^−1^), large multilamellar structures are formed *via* membrane budding from lipid droplets (Figure 1c-d). Notably, when a lower lipid concentration was employed (10 mM), we observed smaller fatty acid droplets and multilamellar vesicles, after heating and cooling ramps, respectively (Movie S2 and Figure S14). Hence, we performed a statistical analysis on the newly-generated multilamellar structures as a function of initial lipid concentration and vesicle size, and incubation time at high temperature (Figure 1e-f and S15). Our data show that higher lipid concentrations and longer incubation times at high temperature result in larger fatty acid droplets – when fatty acid vesicles are only briefly exposed to heat, small lipid droplets do not have enough time to coalesce and thus generate small multilamellar structures upon cooling. At low temperatures, more numerous vesicles, larger in size, can be generated only if lipid droplets are provided sufficient time for efficient membrane budding and self-assembly (Figure S16). Interestingly, the initial diameter of vesicles does not play a role in fatty acid droplet formation or size, and thus has no effect on the size or number of daughter myristoleic acid vesicles (Figure S17). Similar results were obtained for decanoic acid-based vesicles (Figure S18). Overall, our results demonstrate that reversible and tunable pH-induced fatty acid phase transitions, driven by thermal oscillations, support the disassembly and re-assembly of lipid vesicles of varied size and quantity. A critical question to be answered is whether such a simple and robust phenomenon, likely to have occurred recursively and under relatively mild conditions on early Earth, may have supported the onset of a primitive “cell cycle”.

### Release and re-encapsulation of the protocellular content

The thermally-induced vesicle-to-lipid droplet phase transition observed herein explains why fatty acid protocells do not retain compartmentalised genetic material when exposed to high temperatures,^12^ as demonstrated by the specular features of turbidity and leakage profiles (Figure S19). We thus investigated whether the next-generation population of vesicles, formed upon cooling *via* membrane budding from lipid droplets, could re-encapsulate the released genetic content. Myristoleic acid vesicles containing fluorescently-labelled dextran (5kDa), chosen as a model cargo, were either kept at room temperature or subject to a full thermal cycle (25°C→95°C→25°C, at a rate of 0.1°C·s^−1^), then loaded onto a size-exclusion column to evaluate the loss of compartmentalised material (Figure 2a). While most of the fluorescent content (96.8%) was released from vesicles upon heat exposure as previously reported,^12^ a peak corresponding to compartmentalised dextran (3.2%) could be detected at the end of the thermal cycle. There are two possible explanations for such a finding: i) a small fraction of the fatty acid vesicles is not destroyed by the thermal cycle, or ii) the fluorescent content is entirely released during the heating ramp and partially re-encapsulated during the re-assembly of membranes upon cooling. Scenario ii) was confirmed by running the experiment with initially empty vesicles and unencapsulated fluorescently-labelled dextran (Figure 2b). Here, dextran-loaded vesicles were observed only after the thermal cycle, demonstrating that encapsulation takes place during vesicle regeneration.

**Figure 2.**
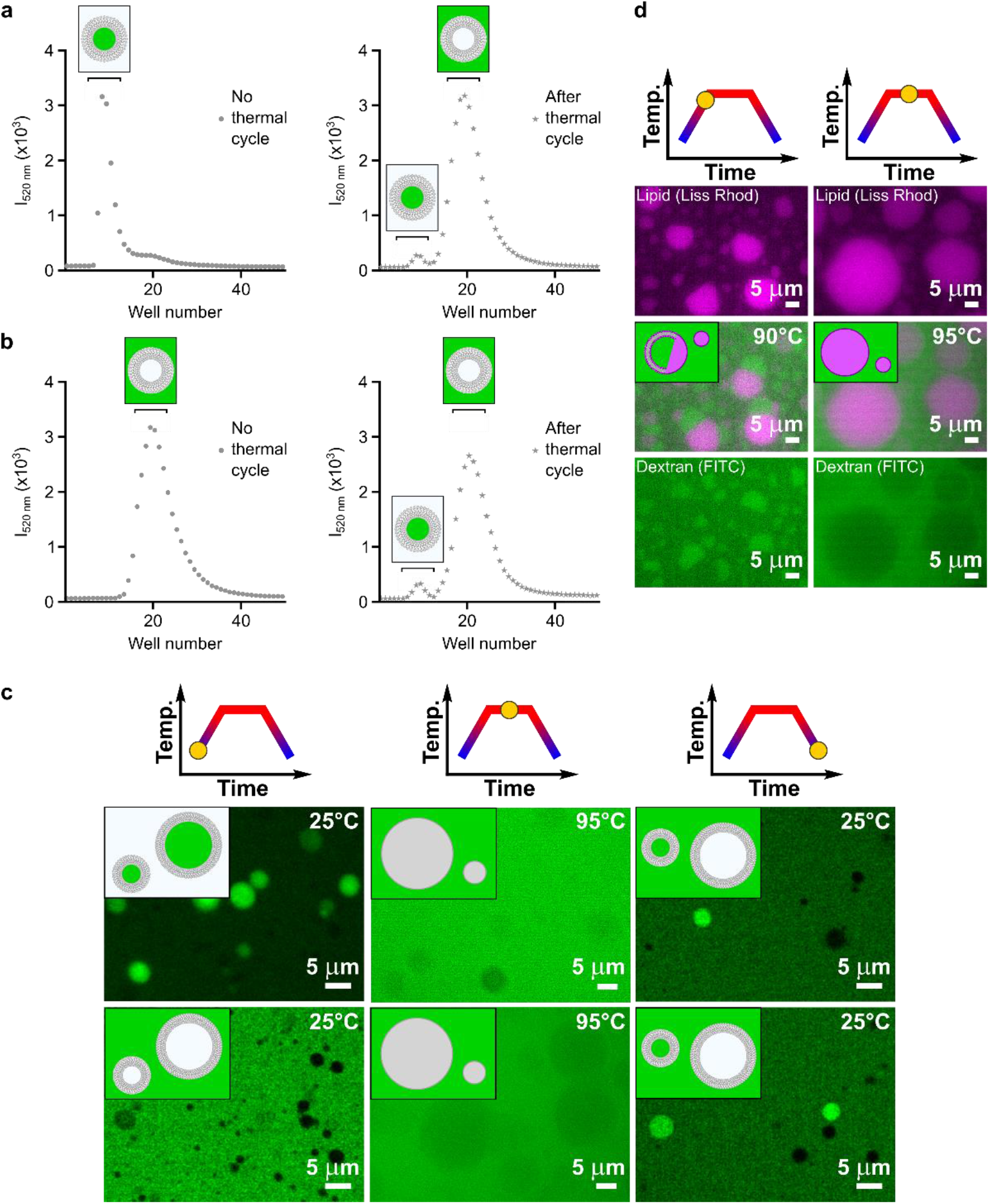
Thermal cycling drives release and re-encapsulation of protocellular content. a) Size-exclusion chromatograms showing partial re-encapsulation (3.2%) of FITC-dextran in 100 nm-radius vesicles made of 50 mM myristoleic acid in 200 mM Tris-HCl, pH 8, upon heat exposure. Content uptake was monitored by fluorescence (λ_exc_ = 495 nm). b) Size-exclusion chromatograms showing partial encapsulation (4.1%) of FITC-dextran in 100 nm-radius vesicles made of 50 mM myristoleic acid in 200 mM Tris-HCl, pH 8, upon heat exposure. Content uptake was monitored by fluorescence (λ_exc_ = 495 nm). c) Microscopy images corresponding to experiments reported in a) (top row) and b) (bottom row) before the thermal cycles (25°C, confocal microscopy images), after the heating ramps (95°C, hot-stage epifluorescence microscopy images) and after the cooling ramps (25°C, confocal microscopy images) are shown for 50 mM myristoleic acid vesicles in 200 mM Tris-HCl, pH 8. d) Hot-stage epifluorescence microscopy images corresponding to the experiment reported in a) at high temperature values (90°C, left; 95°C, right) show the formation of faceted myristoleic acid structures (90°C, left), which transiently trap the aqueous content. Such multilayer structures then convert to lipid droplets (95°C, right), completely releasing the encapsulated material. Data are mean and SEM, n = 3 replicates.

To gain biophysical insight on the lipid droplet-to-vesicle phase transition and the related content re-encapsulation, we repeated both experiments, visualising the process by confocal and hot-stage epifluorescence microscopy (Figure 2c). The mechanism we are proposing is that, when the solution becomes too acidic, unilamellar vesicles begin to collapse at first into multilamellar structures still capable of hosting aqueous milieus, and later into highly dense lipid droplets which expel the protocellular content to the bulk.^28–30^ The observation of faceted structures at high temperatures suggests that the encapsulated fluorescent content is not immediately released in solution, but rather transiently trapped by (multiple) lipid bilayers (Figure 2d). Upon further heat exposure or temperature increase, the encapsulated material is completely excluded, and lipid droplets appear as dark circular spots in the fluorescent background. Along the cooling ramp, as the pH value becomes more suitable for stabilising fatty acid vesicles, the lipid molecules that are present at the surface of the fatty acid droplets begin to self-organise into multilamellar structures (Figure S20) with intercalated fluorescent aqueous content. Thermally-induced pH fluctuations are thus responsible not only for the disassembly and re-assembly of primitive compartments, but also for both the release and subsequent re-uptake of protocellular material.

### Content mixing and reshuffling: a new cohort of protocells

One could imagine that early Earth was inhabited by different populations of protocells, each hosting their own prebiotic content, including genetic material. Those protocells would have likely been exposed to naturally-occurring pH and temperature gradients^19^ and, when made of short-to-medium chain fatty acids, would have undergone the phase transitions discussed in the previous sections. Crucially, when two distinct fatty acid protocell populations undergo such a phase transition, both their encapsulated material and lipids can get mixed by virtue of the assembly/disassembly cycles of their building blocks. Therefore, we propose thermal cycling as a means to potentially generate a new population of protocells with reshuffled content and bilayer components. To investigate this hypothesis, we prepared empty fatty acid protocells and exposed them to three sequential thermal cycles, adding a different fluorescent oligonucleotide to the solution before every cycle. FITC-, Cy3-, and Cy5.5-labelled 10-nucleotide oligomers were selected as proxy for protocellular genetic content. Fluorescence profiles recorded after size-exclusion column chromatography suggest that fluorescent oligonucleotides are re-encapsulated in the second-, third- and fourth-generation of daughter protocells, respectively (Figure S21). Such result further demonstrates the reversibility of membrane assembly and disassembly, as well as content release and uptake, upon recurring thermal cycling. Next, we prepared two distinct vesicle populations, each loaded with a different fluorophore, and mixed them prior to incubation at high temperature; newly-generated vesicles exhibited both fluorescence signals, confirming that the encapsulation of reshuffled content took place (Figure S22). Confocal microscopy images confirm that both fluorescent oligonucleotides are co-compartmentalised within the same protocells (Figure 3a). As thermally-driven phase transitions yield new populations of daughter fatty acid protocells, we sought to explore whether new blended membranes would also be generated by lipid mixing in the oil phase. Large mixed multilamellar structures were observed upon heating from combinations of fluorescently-labelled unilamellar protocells (Figure 3b). No content nor lipid reshuffling could be detected for populations of vesicles only mixed after thermal cycling (Figure S23-24).

**Figure 3.**
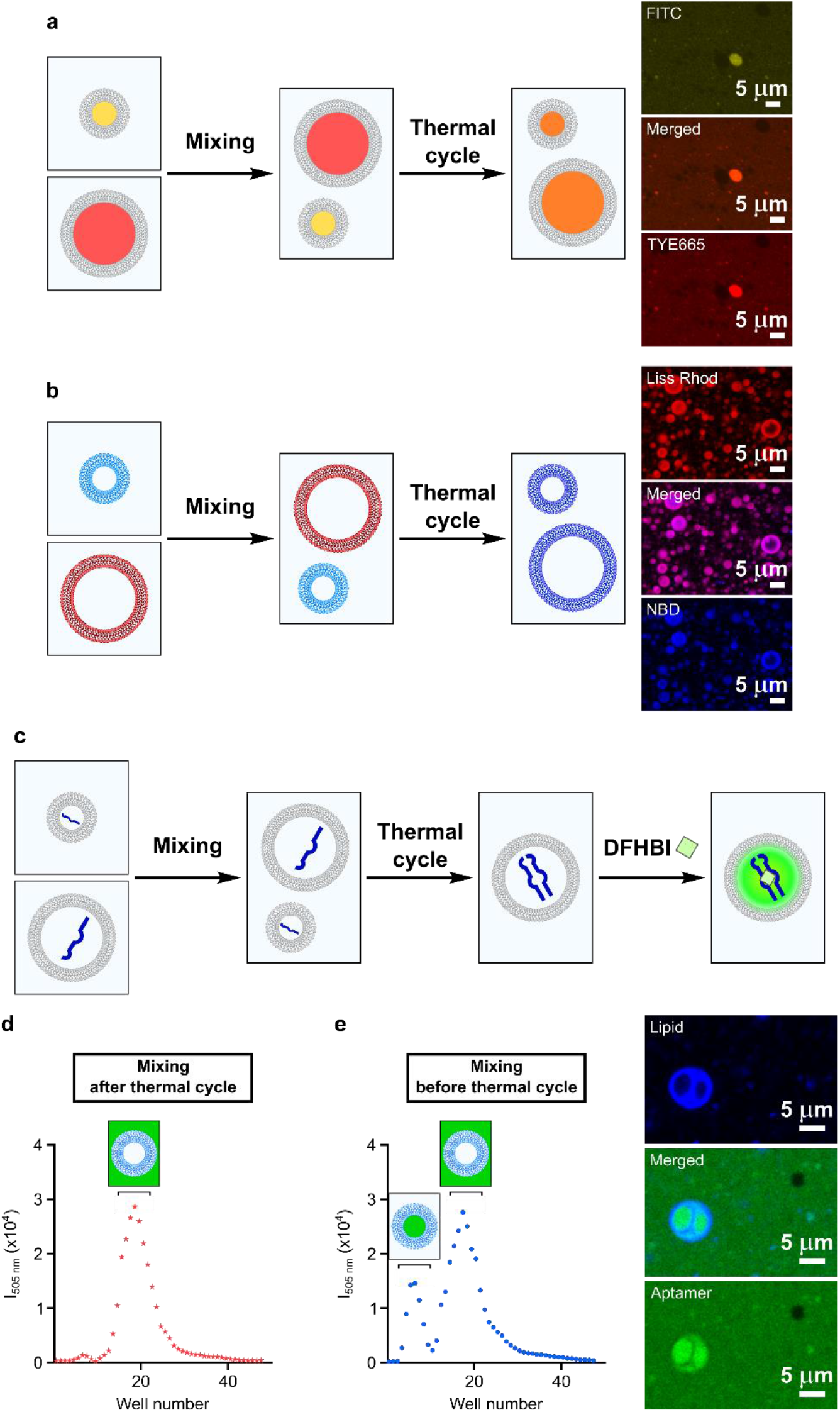
Thermal cycling drives reassembly of protocells with reshuffled membrane material and encapsulated content. a) Schematic representation and confocal microscopy images for experiments with populations of vesicles with different fluorescent content. 100 nm-radius vesicles, made of 50 mM myristoleic acid in 200 mM Tris-HCl, pH 8 and containing either FITC-10nt or TYE665-10nt oligonucleotides, were mixed before undergoing thermal cycling and, after 1 h re-equilibration at room temperature, were visualised by confocal microscopy. b) Schematic representation and confocal microscopy images for experiments with populations of vesicles, labelled with different fluorescent lipids. 100 nm-radius vesicles, made of 50 mM myristoleic acid in 200 mM Tris-HCl, pH 8 and either NBD-PE or Rh-DHPE, were mixed before undergoing thermal cycling and, after 1 h re-equilibration at room temperature, were visualised by confocal microscopy. c) Schematic representation of the experiment to reconstitute a split Broccoli aptamer inside myristoleic acid vesicles. d) Size-exclusion chromatograms showing no reconstitution of the split Broccoli aptamer when vesicle mixing occurs after thermal cycling. e) Size-exclusion chromatograms showing reconstitution of the split Broccoli aptamer (20.3%) when vesicle mixing occurs before thermal cycling. Experiments for Broccoli aptamer reconstitution were performed with 100 nm-radius vesicles made of 50 mM myristoleic acid in 200 mM Tris-HCl, pH 8. Broccoli aptamer reconstitution was monitored by fluorescence of DFHBI (λ_exc_ = 505 nm). Confocal images were collected prior to purification. Data are mean and SEM, n = 3 replicates.

An interesting advantage of protocellular reshuffling is the potential for a new cohort of primitive compartments to arise with novel (genetic) functionalities. For example, combining genetic material that is independently synthesised within distinct fatty acid vesicles could give rise to new rounds of nucleic acid polymerisation, as well as ribozyme-based catalytic processes. As a proof-of-concept, we designed a set of experiments aimed at reconstituting a shorter split version of the Broccoli aptamer within the new population of protocells, generated *via* thermal cycling (Figure 3c). When two populations of vesicles are first exposed to thermal ramps and later mixed, no fluorescence from the encapsulated DFHBI fluorogen can be detected (Figure 3d). However, when a mixed sample of different protocells undergoes thermal cycling, the newly generated vesicles incorporate both RNA strands, thus efficiently assembling the Broccoli aptamer, which enhances the fluorescent signal of the DFHBI fluorogen (Figure 3e). Together, these results provide strong evidence that thermally-driven pH oscillations can drive reversible and tunable lipid phase transitions and support the release and reshuffle of protocellular genetic components, thus leading to the emergence of a new generation of protocells with potentially enhanced functionalities.

## Discussion

In the absence of modern biological machinery, primitive cells likely had to rely on the self-assembling and dynamic properties of their components and on interactions with their environment to achieve basic cellular functions, including primitive cellular replication. Primitive cell cycles involving alternating dehydration and rehydration steps have been recently proposed;^32^ however, volatile reactants can easily escape and pH- or temperature-sensitive substrates can undergo irreversible degradation during dehydration. Alternatively, primitive cell cycles could be based on recursive growth and division of lipid vesicles^10^. Despite better mimicking modern cellular pathways, such iterations would not support the uptake, release and exchange of protocellular content with the environment.

In this context, we propose a primitive cell cycle based on mild, naturally-occurring environmental fluctuations, which support alternating membrane dissolution and reorganisation steps. The pathway for protocell disassembly and reassembly described herein is robust, in that it operates in different membrane compositions, over a wide range of buffered and unbuffered solutions, as well as in the presence of salts, nucleotides and short peptides. Temperature-driven pH oscillations result in reversible fatty acid phase transitions, which enable the release of products (which might have feedback-inhibited further rounds of compartmentalised prebiotic reactions), the uptake of membrane-impermeable substrates (which might have been required for both anabolic and catabolic pathways) and the exchange of material between different protocells (which might have driven selection processes), as well as the reshuffling of lipophilic components (which might have yielded more modern-like membranes and/or membranes with superior traits, such as improved stability and selective permeability).

Before the evolution of modern lipids and transport systems, primitive cells may have depended on simple physical processes for their replication. In our novel model, we innovatively propose to take advantage of the instability of fatty acids in order to drive recurrent cycles of protocell disassembly and reassembly, in which both the parental membrane material and encapsulated content are transmitted to a new cohort of daughter protocells. Only the iterative temperature-driven reshuffling of lipid and genetic material could have then led to a cascade of new selective pressures for the evolution of more advanced protocells to overcome the intrinsic instability of prebiotic lipids.^33^

If such a primitive cell cycle could be coupled with nonenzymatic nucleic acid replication, it would set the stage for functional nucleic acids to competitively emerge and support early stages of Darwinian evolution. Additionally, our discovery might hint at a prebiotic means of recycling materials of metabolically “dead” protocells. Hence, the identification of prebiotically-relevant primitive cell cycles like the one proposed herein, supported by mild environmental fluctuations, may represent a step towards the emergence of advanced protocells, which can help in designing artificial systems with life-like behaviours and potentially gaining greater insight into the origin of modern cells.

## Acknowledgements

The authors gratefully acknowledge support from the European Union’s Horizon 2020 research and innovation programme under the Marie Skłodowska-Curie (RNA-Rep, MSCA grant no. 839899 to C.B.) and the European Research Council (NANOCELL, ERC-STG no. 851667 to L.D.M.), the Mexican National Council for Science and Technology (CONACYT, grant no. 472427 to R.R.S.), Cambridge Trust (to R.R.S.), the EPSRC CDT in Nanoscience and Nanotechnology (NanoDTC, grant no. EP/L015978/1 to R.R.S.), the Natural Sciences and Engineering Research Council of Canada (NSERC, Early Career Researcher Grant no. 401667 to D.K.O.) and the Royal Society (University Research Fellowship grant no. UF160152 to L.D.M.). The authors thank Lesley McKeane, MRC LMB Visual Aids Department, for support with graphics and movies, and the MRC LMB Electron Microscopy Facility for access and support of electron microscopy sample preparation and data collection. The authors acknowledge Dr. Gianluca Petris, Dr. David Russell and Prof. Jack Szostak for fruitful discussions.

## Data availability

The authors declare that the data supporting the findings of this study are available from the authors upon reasonable request.

