## Supporting information for "Thermal oscillations enable reshuffling of genetic material in a primitive cell cycle"

**Materials and methods**

**Supporting Figures 1 to 24**

**Supporting References**

### Materials and methods

Reagents and solvents were bought from Nu-Chek Prep, Avanti Polar Lipids, Hellma Analytics, CM Scientific, AMS Technologies, Merck and Thermo Fisher and were used without further purification unless otherwise stated. Oligonucleotides were purchased from IDT. For microscopy studies, the fluorescent phospholipids N-(7-nitrobenz-2-oxa-1,3-diazol-4-yl)-1,2-dihexadecanoyl-sn-glycero-3-phosphoethanolamine (NBD PE) and Lissamine rhodamine B 1,2-dihexadecanoyl-sn-glycero-3-phosphoethanolamine (Rh-DHPE) were used (Life Technologies).

A Mettler Toledo SevenEasy pH Meter S20 was used to monitor the pH of the solutions, adjusted with either NaOH or HCl solutions as appropriate. UV-vis analyses were performed on a Varian Cary 6000i UV/Vis/NIR spectrophotometer and fluorescence measurements were recorded on a Varian Cary Eclipse fluorescence spectrophotometer, both equipped with a multi-sample Peltier temperature controller. For confocal microscopy, vesicles were imaged on a Zeiss LSM 780 confocal microscope equipped with a 63× oil objective. Vesicles were purified by size-exclusion column chromatography (Sepharose 4B); fractions were collected in 96-well plates (Costar, black, clear bottom) using a fraction collector FC203B (Gilson) and analysed with a PHERAstar FSX plate reader.

Peak integration on chromatograms was performed using Origins (version 2019). Statistical analysis (ordinary one-way ANOVA and simple linear regression) was performed using GraphPad Prism (version 9.0.1). Images were processed with Fiji[1]. All data shown are representative of distinct samples, with n = 3 replicates. Unless otherwise stated, data are mean and SEM.

**Abbreviations**. The following abbreviations are used throughout: DA, decanoic acid; DOH, decanol; MA, myristoleic acid.

**Modelling**. The software package HoloPy[2] was used to calculate the exact Lorenz-Mie solution for how various vesicle samples scatter light[3]. Parameters of 2 mm sample path length, 420 nm illumination light, 100 nm extruded vesicle radius, 2 nm lipid bilayer thickness[4], 0.2 $nm^2$ average area per lipid[5] and 5.2 mM critical vesicle concentration were used for modelling.

**Cryogenic electron microscopy (cryo-EM)**. Vesicle samples were applied to freshly glow-discharged (40 mA, 1 min) EM holey grids (Quantifoil Cu/Rh R22 200 mesh) and plunge-frozen in liquid ethane using a Vitrobot Mark IV (Thermo Fisher), with blot force -15, blot time 2.5 s, humidity 100% and at 4°C. The grids were imaged with a Glacios (Thermo Fisher) electron microscope operated at 200 kV at the nominal magnification of 57,000×, corresponding to a pixel size of 2.55 Å. Images were acquired on a Falcon 3 detector (Thermo Fisher) in linear mode and with a total dose of $60e^{-}\cdot Å^{-2}$.

**Hot stage epifluorescence microscopy.** Borosilicate glass capillaries (internal section of 2 × 0.2 mm) were cleaned through sonication cycles of 15 minutes in Hellmanax III 2%/Isopropanol/MilliQ water. Once dried, one end of the capillary was sealed with optical glue, cured under UV light (λ = 365 nm) for 5 minutes. The capillaries were subsequently filled with fluorescently-labelled vesicle samples up to approximately 85% of their volume. The remainder was filled with mineral oil. Afterwards, the capillary was stuck to a glass coverslip using a two-part epoxy resin and hardener. Glass capillaries on coverslips were placed on a copper plate connected to a Peltier element and fixed with aluminium tape. This set-up enabled fine control over temperature. Imaging was done using a home-built Nikon Eclipse Ti-E inverted microscope, equipped with a 40× objective lens (Nikon, Plan APO λ, N.A. 0.95) and a Grasshopper3 GS3-U3-23S6M camera (Point Gray Research). The illumination was provided by single-colour light emitting diodes (LEDs) using a filter set for TexasRed or Green Fluorescent Protein. Temperature ramps were performed using a custom-built script, enabling precise manipulation of the instrument in terms of time, temperature, and illumination as required.

**Preparation of vesicles.** Vesicles made of myristoleic acid or decanoic acid:decanol (2:1 ratio) were prepared by direct dispersion in an aqueous buffered solution as previously reported.[6] Samples were briefly vortexed, tumbled at room temperature for 30 min and then extruded with 11 passages through a 100 nm pore membrane (Whatman) using a Mini-Extruder (Avanti Polar Lipids). To encapsulate fluorescent material, 1 mM FITC- or Texas Red-labelled dextran (mol wt 4000 Da) or 10 μM labelled 10-nt oligonucleotides (5′-TGTGCCAGTA-3′, fluorophores: FITC, Cy3, Cy5.5, TYE665) was added to the buffered solution prior to resuspension. When loaded with fluorescent cargo, vesicles were loaded onto a size-exclusion column (Sepharose 4B, 6 mL) to remove unencapsulated material. Unless otherwise stated, 0.2 M Tris-HCl, pH 8 was chosen as the running buffer, supplemented with 25 mM decanoic acid:decanol (2:1 ratio). Elution was monitored by fluorescence and the fractions containing vesicles were collected. To evaluate encapsulation upon thermal cycling, 10 μM FITC-dextran or labelled oligonucleotides was added to vesicle solutions prior to heat exposure. For microscopy and fluorescence studies, vesicles were prepared by thin film hydration methods[7] on the bottom of glass vials. Vesicles, made of fatty acids, contained 0.15 mol % Liss Rh-PE or NBD-PE (for microscopy) and 0.2 mM Laurdan (for fluorescence). Solvent was evaporated for >12 h, and, unless otherwise stated, lipid films were hydrated with 0.2 M Tris-HCl, pH 8. Alternatively, non-fluorescent vesicles, prepared by direct suspension methods, were stained with Nile Red dye prior to visualisation.

**Thermal cycling.** For turbidity and fluorescence analysis, fatty acid vesicles were heated up to 95°C (at a rate of 0.1°C·s$^{-1}$) and kept at 95°C for 15 minutes, then cooled down to 25°C at the same rate, while recording absorbance values at 420 nm or fluorescence intensity at 425 nm and 495 nm. The same thermal settings were used for hot stage epifluorescence microscopy. For confocal microscopy and cryo-EM, samples were visualised before and after thermal cycles (in the latter case, after waiting 1 hour for re-equilibration of the sample at room temperature).

**Leakage studies.** Vesicles containing 1 mM FITC-dextran were prepared, extruded and purified as previously described. The purified vesicles were incubated at a constant temperature (20-95°C) for 1 h. Vesicles were then purified, and fractions were analysed by fluorescence spectrophotometry ($\lambda_{exc}$ = 454 nm). The retention percentage of FITC-dextran was reported as a function of temperature.

**Reconstitution of a split Broccoli aptamer inside vesicles.** A minimal version of the Broccoli aptamer was split into two oligonucleotides (a 23-nt oligonucleotide (5′-GCGGAGACGGUCGGGUCCAGAUA-3′) and a 28-nt oligonucleotide (5′-UAUCUGUCGAGUAGAGUGUGGGCUCCGC-3′)). Two different batches of vesicles, made of myristoleic acid, were prepared by direct dispersion in an aqueous buffered solution, containing 100 μM of each oligonucleotide. Vesicles were extruded and purified as previously described. Purified vesicles, containing different oligonucleotides, were mixed either before or after heat exposure and purified again, all fractions being collected in 96 well plates. A solution containing DFHBI (stock solution: 20 mg/mL in DMSO) and KCl was prepared and added to each well (final concentrations: 2 μM DFHBI and 100 mM KCl). Plates were left at 25°C for 2 hours to account for DFHBI equilibration across fatty acid membranes. Fluorescence was then recorded between 475 and 575 nm (excitation wavelength: 447 nm) and vesicles were visualised with confocal microscopy.

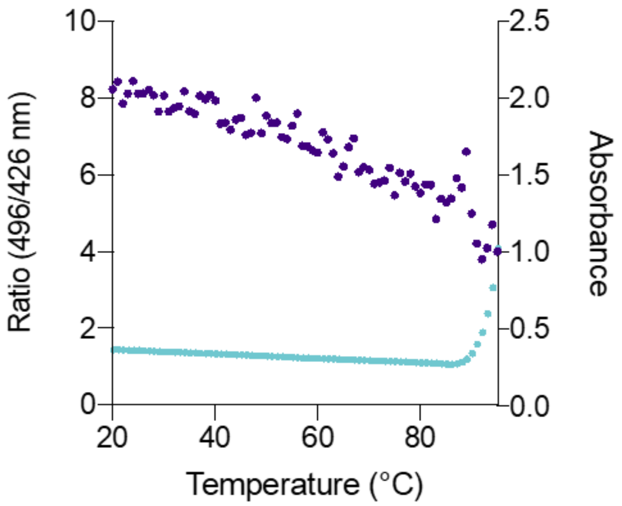


*Figure S1 – Overlapped profiles of turbidity and fluorescence intensity as a function of temperature for 100 nm-radius vesicles made of 50 mM myristoleic acid (in 200 mM Tris-HCl, pH 8). Absorbance (light turquoise) is monitored at 420 nm, whereas Laurdan fluorescence (purple) is monitored at $\lambda_{exc}$ = 426 and 496 nm. An increase in turbidity and a decrease in fluorescence ratio can be observed above 90°C. n = 3 replicates.*

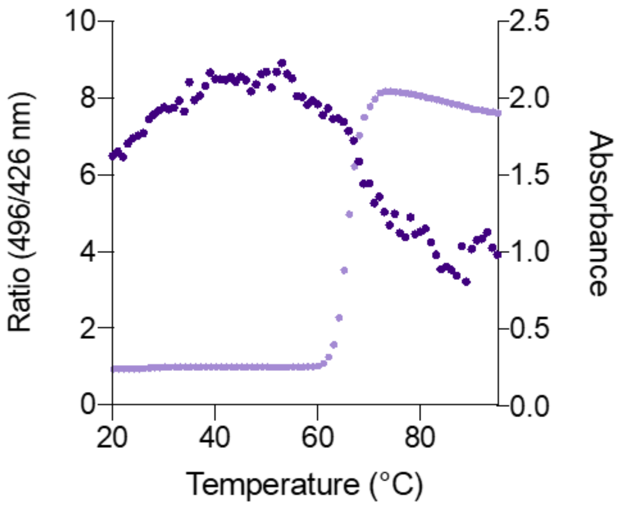


*Figure S2 – Overlapped profiles of turbidity and fluorescence intensity as a function of temperature for 100 nm-radius vesicles made of 50 mM decanoic acid:decanol (2:1 ratio, in 200 mM Tris-HCl, pH 8). Absorbance (light violet) is monitored at 420 nm, whereas Laurdan fluorescence (purple) is monitored at $\lambda_{exc}$ = 426 and 496 nm. An increase in turbidity and a decrease in fluorescence ratio can be observed around 65°C. n = 3 replicates.*

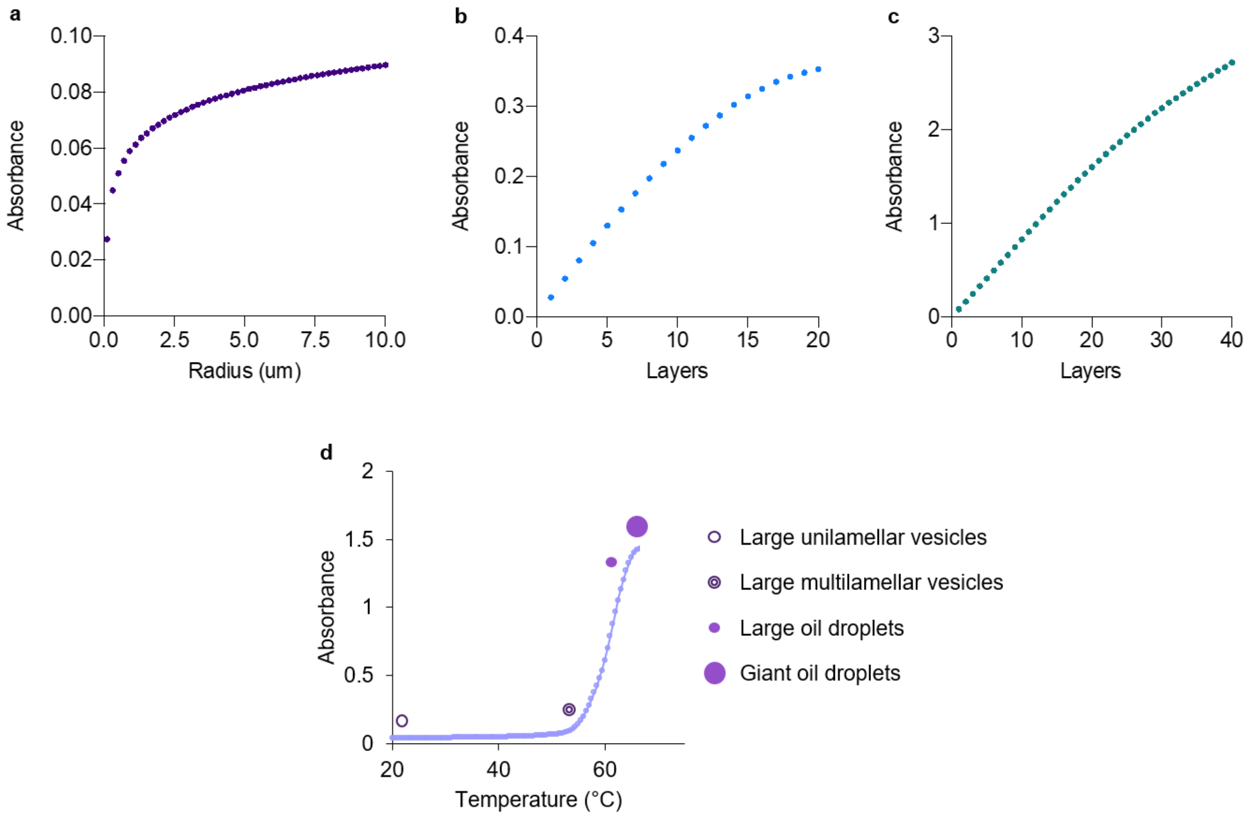


*Figure S3 – Modelling data for decanoic acid-based vesicles. Absorbance data are modelled for 25 mM decanoic acid:decanol (2:1 ratio). a) Modelled absorbance for a sample made of unilamellar vesicles as a function of radius. b) Modelled absorbance for a sample made of 100 nm-radius vesicles as a function of lamellarity. c) Modelled absorbance for a sample made of 5 µm-radius sample as a function of lamellarity. d) Experimental turbidity profile with different lipid structures sketched alongside for comparison according to calculated absorbance values.*

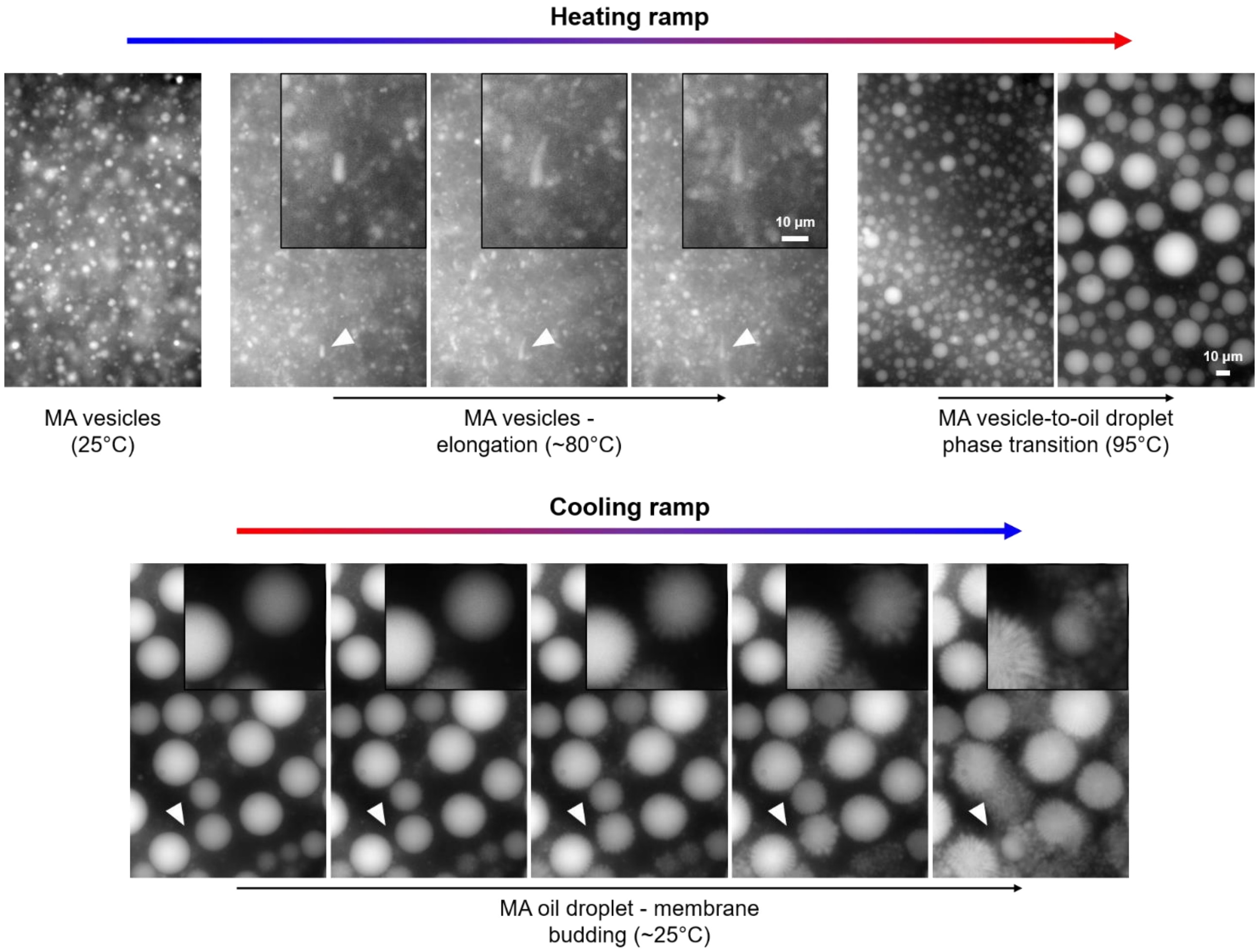


*Figure S4 – Frames extracted from movie S1, recorded during hot stage epifluorescence microscopy, for myristoleic acid vesicles. Before reaching the phase transition temperature (~85°C), myristoleic acid vesicles (50 mM) are first subject to elongation. Above the phase transition temperature, vesicles collapse into small oil droplets, which coalesce while kept at high temperature (top). During the cooling ramp, membrane budding can be observed from the surface of oil droplets (bottom).*

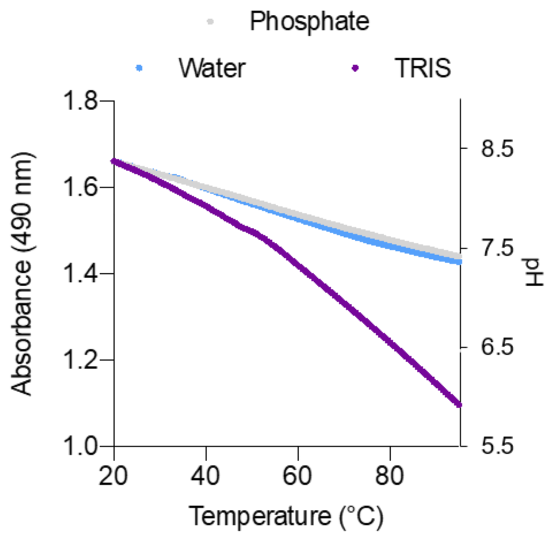


*Figure S5 – Absorbance profiles for fluorescein as a function of temperature in different buffers. Fluorescein was selected as a probe for its pH sensitivity. Phosphate, Tris-HCl and water were heated up to 95°C (at a rate of 0.1°C·s$^{-1}$) in the presence of 10 μM fluorescein and absorbance was monitored at 490 nm. n = 3 replicates.*

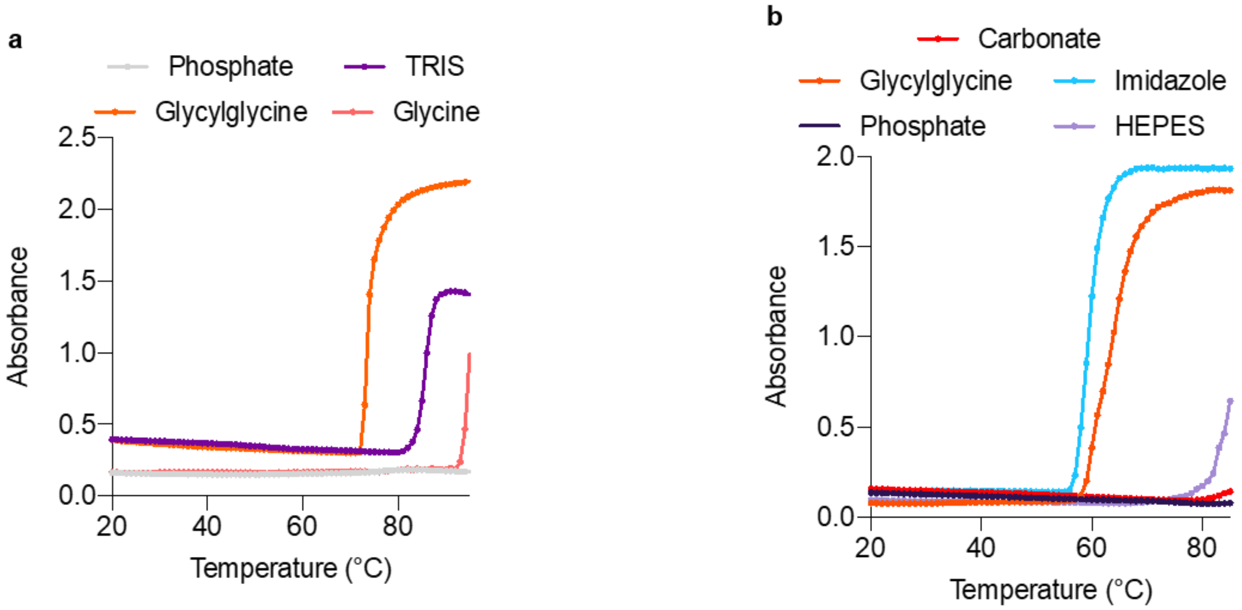


*Figure S6 – Turbidity profiles for fatty acid vesicles in different buffers as a function of temperature. a) Turbidity profile for 100 nm-radius vesicles made of 50 mM myristoleic acid in 200 mM buffer (Tris-HCl, glycylglycine, glycine and phosphate), pH 8. Absorbance was monitored at 420 nm. b) Turbidity profile for 100 nm-radius vesicles made of 50 mM decanoic acid:decanol (2:1 ratio) in 200 mM buffer (Tris-HCl, glycylglycine, imidazole, carbonate and phosphate), pH 8. Absorbance was monitored at 420 nm. A sharp increase in turbidity is observed at different temperatures for different buffers. n = 3 replicates.*

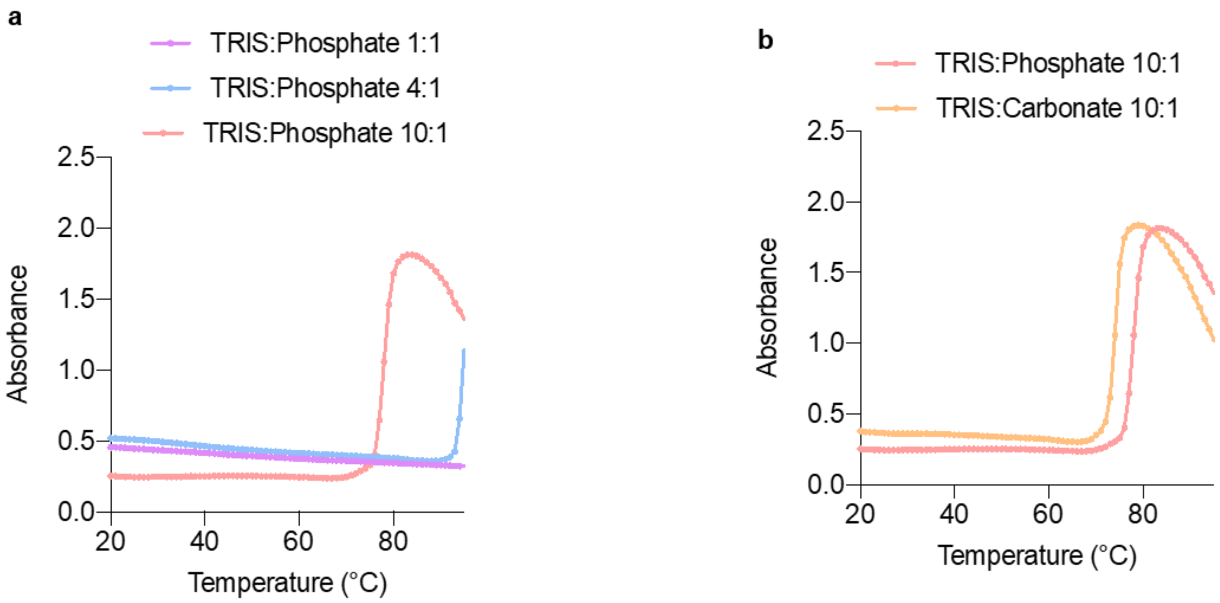


*Figure S7 – Turbidity profiles for myristoleic acid vesicles in different buffer mixtures as a function of temperature. a) Turbidity profile for 100 nm-radius vesicles made of 50 mM myristoleic acid in 200 mM mixed Tris-HCl:phosphate buffer (1:1, 4:1 or 10:1 ratio), pH 8. Absorbance was monitored at 420 nm. b) Turbidity profile for 100 nm-radius vesicles made of 50 mM myristoleic acid in 200 mM mixed Tris-HCl:X buffer, 10:1 ratio (X = phosphate or carbonate), pH 8. Absorbance was monitored at 420 nm. Even though a phase transition is not observed for myristoleic acid vesicles in pure phosphate and carbonate buffers (due to their $pK_a$ being independent of temperature), a sharp increase in turbidity is observed when Tris-HCl is added to either phosphate or carbonate buffers. n = 3 replicates.*

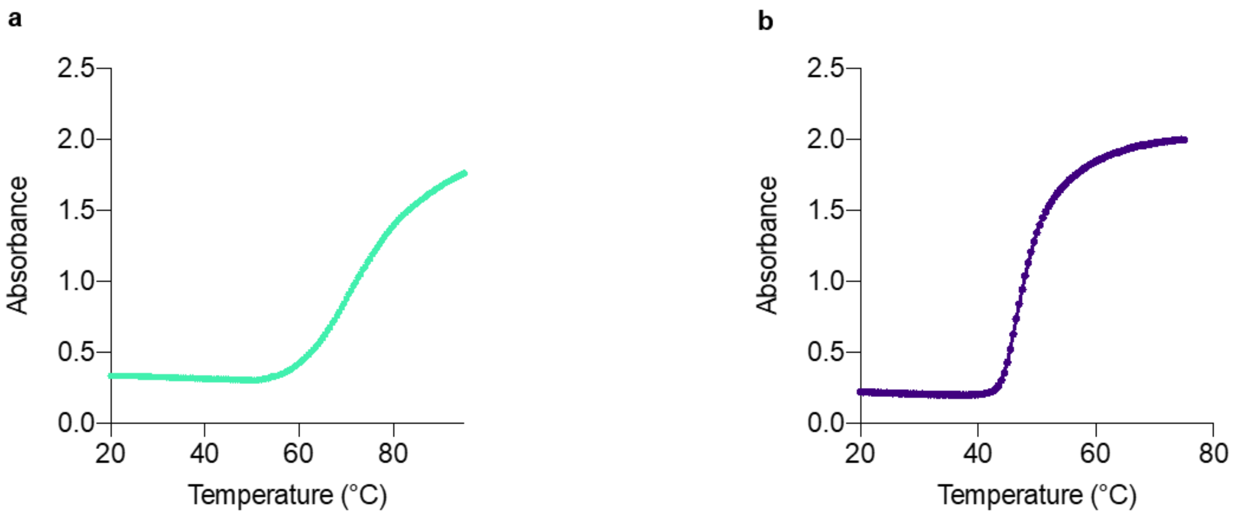


*Figure S8 – Turbidity profiles for fatty acid vesicles in self-buffering conditions as a function of temperature. a) Turbidity profile for 100 nm-radius vesicles made of 50 mM myristoleic acid in water, pH 8. Absorbance was monitored at 420 nm. b) Turbidity profile for 100 nm-radius vesicles made of 50 mM decanoic acid:decanol (2:1 ratio) in water, pH 8. Absorbance was monitored at 420 nm. An increase in turbidity is observed for both fatty acid systems at lower temperature values compared to the results observed in buffered solutions. n = 3 replicates.*

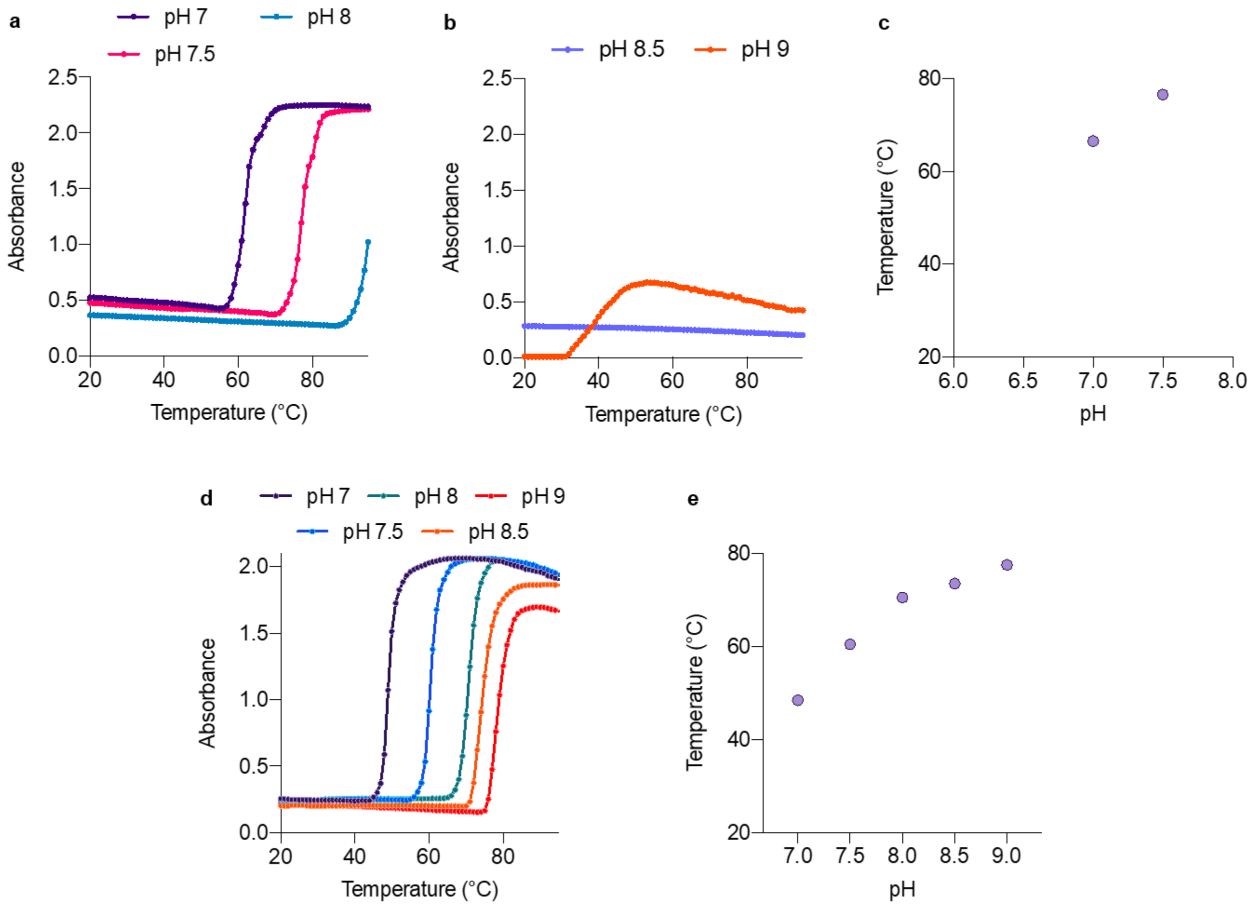


*Figure S9 – Turbidity profiles for fatty acid vesicles made in different pH conditions as a function of temperature. a) Turbidity profile for 100 nm-radius vesicles made of 50 mM myristoleic acid in 200 mM Tris-HCl, at different initial pH values (7, 7.5 and 8). Absorbance was monitored at 420 nm. The increase in turbidity is observed at higher temperatures for higher initial pH values, as the buffer reaches a pH value lower than 6.5 (when fatty acid vesicles become unstable) at higher temperatures, when vesicles are prepared in 200 mM Tris-HCl buffer at higher pH values. b) Turbidity profile for 100 nm-radius vesicles made of 50 mM myristoleic acid in 200 mM Tris-HCl, at different pH values (8.5 and 9). Absorbance was monitored at 420 nm. No destabilisation can be observed when myristoleic acid vesicles are prepared in 200 mM Tris-HCl buffer at pH 8.5, as the buffer does not reach pH values low enough to destabilise vesicles. When myristoleic acid is hydrated with 200 mM Tris-HCl buffer, pH 9, it self-assembles into myristoleate micelles[8,9] – the increase observed in the turbidity profile is due to the micelle-to-vesicle phase transition. c) Temperature vs pH graph with vesicle-to-oil droplet phase transition values extrapolated from the curves in a). d) Turbidity profile for 100 nm-radius vesicles made of 50 mM decanoic acid:decanol (2:1 ratio) in 200 mM Tris-HCl, at different initial pH values (7, 7.5, 8, 8.5, 9). Absorbance was monitored at 420 nm. The increase in turbidity follows the same trend observed for myristoleic acid vesicles. As decanoic acid-based vesicles are more stable at higher pH values in the presence of decanol, vesicles are formed even at an initial pH value of 9. e) Temperature vs pH graph with vesicle-to-oil droplet phase transition values extrapolated from the curves in d). n = 3 replicates.*

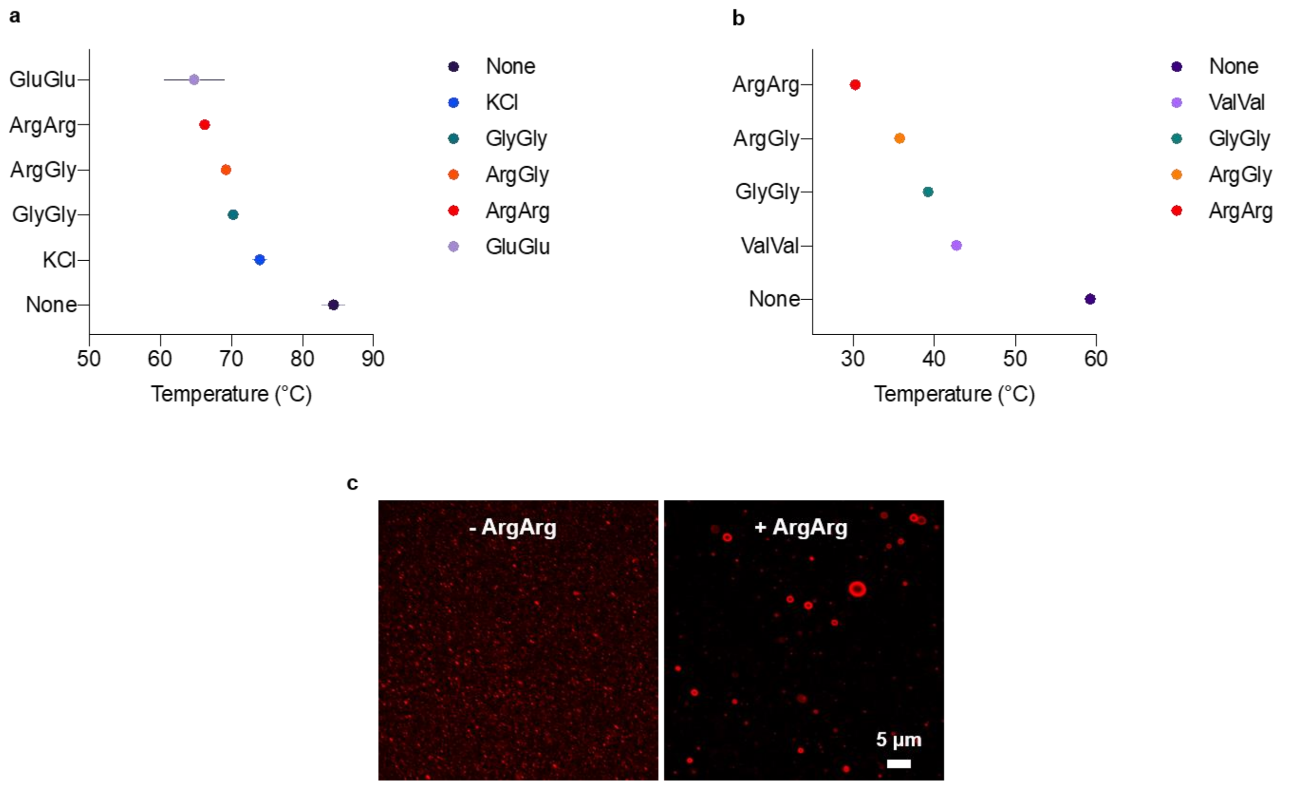


*Figure S10 – Effect of additives on phase transition temperatures for fatty acid vesicles. a) Phase transition temperatures observed for vesicles made of 50 mM myristoleic acid in 200 mM Tris-HCl, pH 8 in the presence of 5 mM additive (salt or peptides). Values have been extrapolated from turbidity curves. b) Phase transition temperatures observed for vesicles made of 50 mM decanoic acid:decanol (2:1 ratio) in 200 mM Tris-HCl, pH 8 in the presence of 5 mM additive (peptides). Values have been extrapolated from turbidity curves. The observed trend suggests that an increase in ionic strength, as well as additional electrostatic interactions can dramatically decrease the phase transition temperature for both fatty acid systems. c) Confocal microscopy images taken for 100 nm-radius vesicles made of 50 mM decanoic acid:decanol (2:1 ratio) in 200 mM Tris-HCl, pH 8 in the presence or absence of arginylarginine (ArgArg), after heating them up to 45°C, and then cooling them down to 25°C and re-equilibrating them for 1 h. In the presence of the peptide, the vesicle-to-oil droplet transition occurs at low temperatures (~30°C); in the absence of the peptide, the phase transition occurs at higher temperatures (~60°C). When the samples are heated up to 45°C, only in the presence of the peptide vesicles undergo phase transition to oil droplets and reassemble, upon cooling, in larger multilamellar vesicles. n = 3 replicates.*

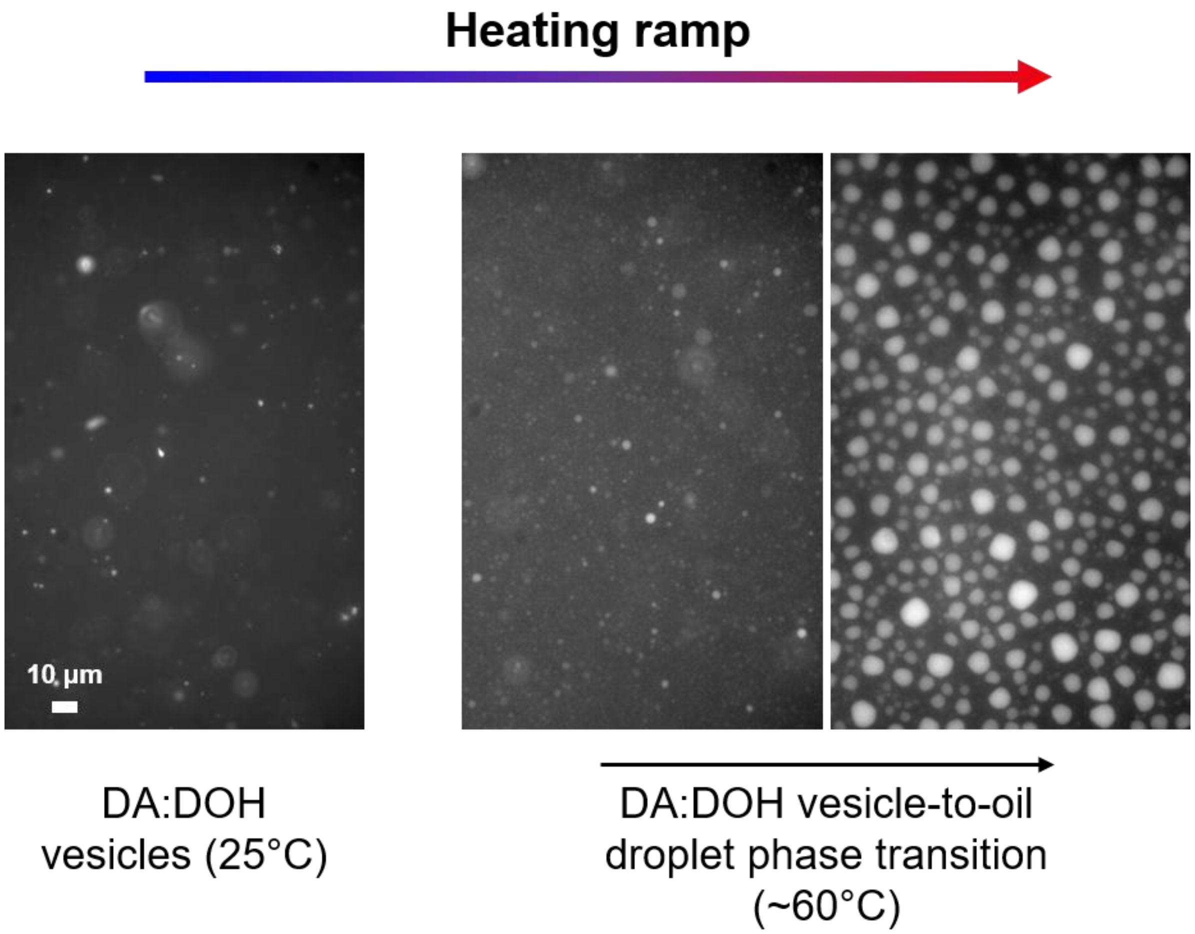


*Figure S11 – Hot stage epifluorescence microscopy images collected for decanoic acid-based vesicles. Once the phase transition temperature is reached (~60°C), decanoic acid:decanol (2:1 ratio) vesicles (50 mM) collapse into small oil droplets, which merge while kept at high temperature. Faceted structures can be observed at high temperatures during oil droplets merging.*

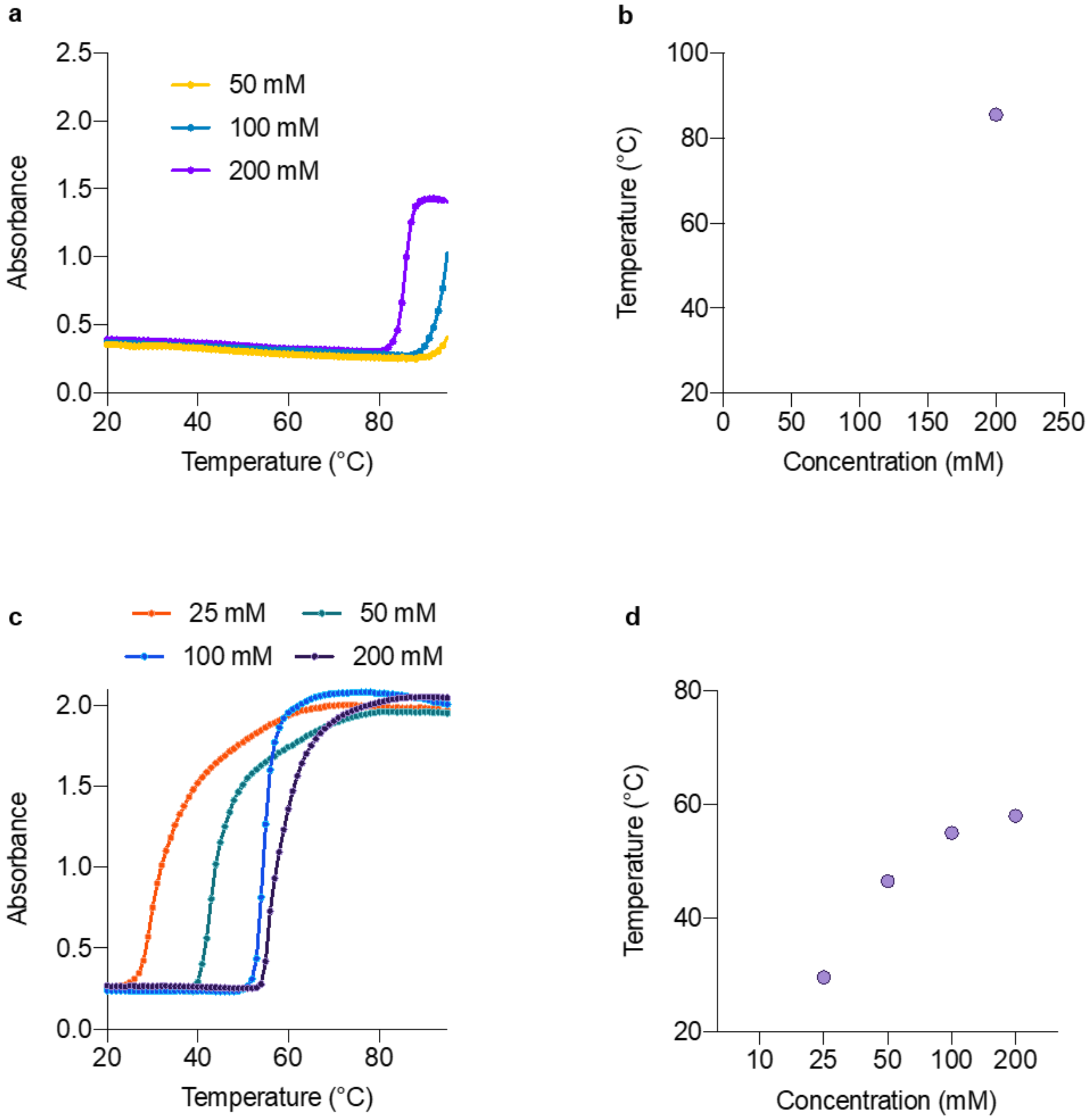


*Figure S12 – Turbidity profiles for fatty acid vesicles made in Tris-HCl (various concentrations) as a function of temperature. a) Turbidity profile for 100 nm-radius vesicles made of 50 mM myristoleic acid in Tris-HCl (200 mM, 100 mM or 50 mM), pH 8. Absorbance was monitored at 420 nm. Phase transition temperature increases for lower buffer concentrations, likely because pH decreases more slowly for lower buffer concentrations. b) Temperature vs pH graph with vesicle-to-oil droplet phase transition values extrapolated from the curves in a). c) Turbidity profile for 100 nm-radius vesicles made of 50 mM decanoic acid:decanol (2:1 ratio) in Tris-HCl (200 mM, 100 mM, 50 mM or 25 mM), at pH 8. Absorbance was monitored at 420 nm. Phase transition temperature decreases for lower buffer concentrations, likely because of the higher instability of decanoic acid-based vesicles at lower buffer concentrations. d) Temperature vs pH graph with vesicle-to-oil droplet phase transition values extrapolated from the curves in c). n = 3 replicates.*

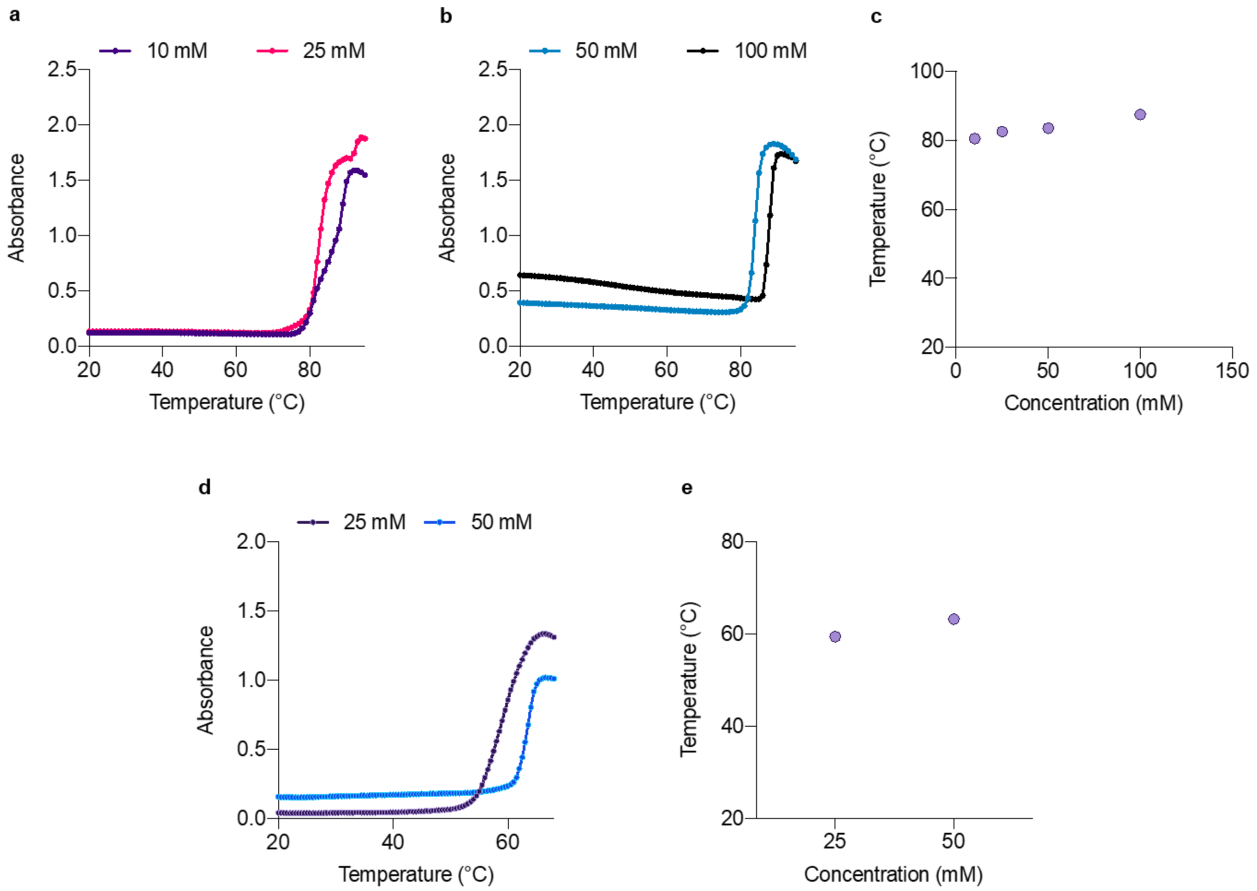


*Figure S13 – Turbidity profiles for fatty acid vesicles made in various fatty acid concentrations as a function of temperature. a) Turbidity profile for 100 nm-radius vesicles made of myristoleic acid (25 mM or 10 mM) in 200 mM Tris-HCl, pH 8. Absorbance was monitored at 420 nm. Further investigations are undergoing to understand the origin of the two inflection points observed in the turbidity profiles. b) Turbidity profile for 100 nm-radius vesicles made of myristoleic acid (100 mM or 50 mM) in 200 mM Tris-HCl, pH 8. Absorbance was monitored at 420 nm. c) Temperature vs pH graph with vesicle-to-oil droplet phase transition values extrapolated from the curves in a) and b). d) Turbidity profile for 100 nm-radius vesicles made of decanoic acid:decanol (2:1 ratio, 50 mM or 25 mM) in 200 mM Tris-HCl, at pH 8. Absorbance was monitored at 420 nm. e) Temperature vs pH graph with vesicle-to-oil droplet phase transition values extrapolated from the curves in d). n = 3 replicates.*

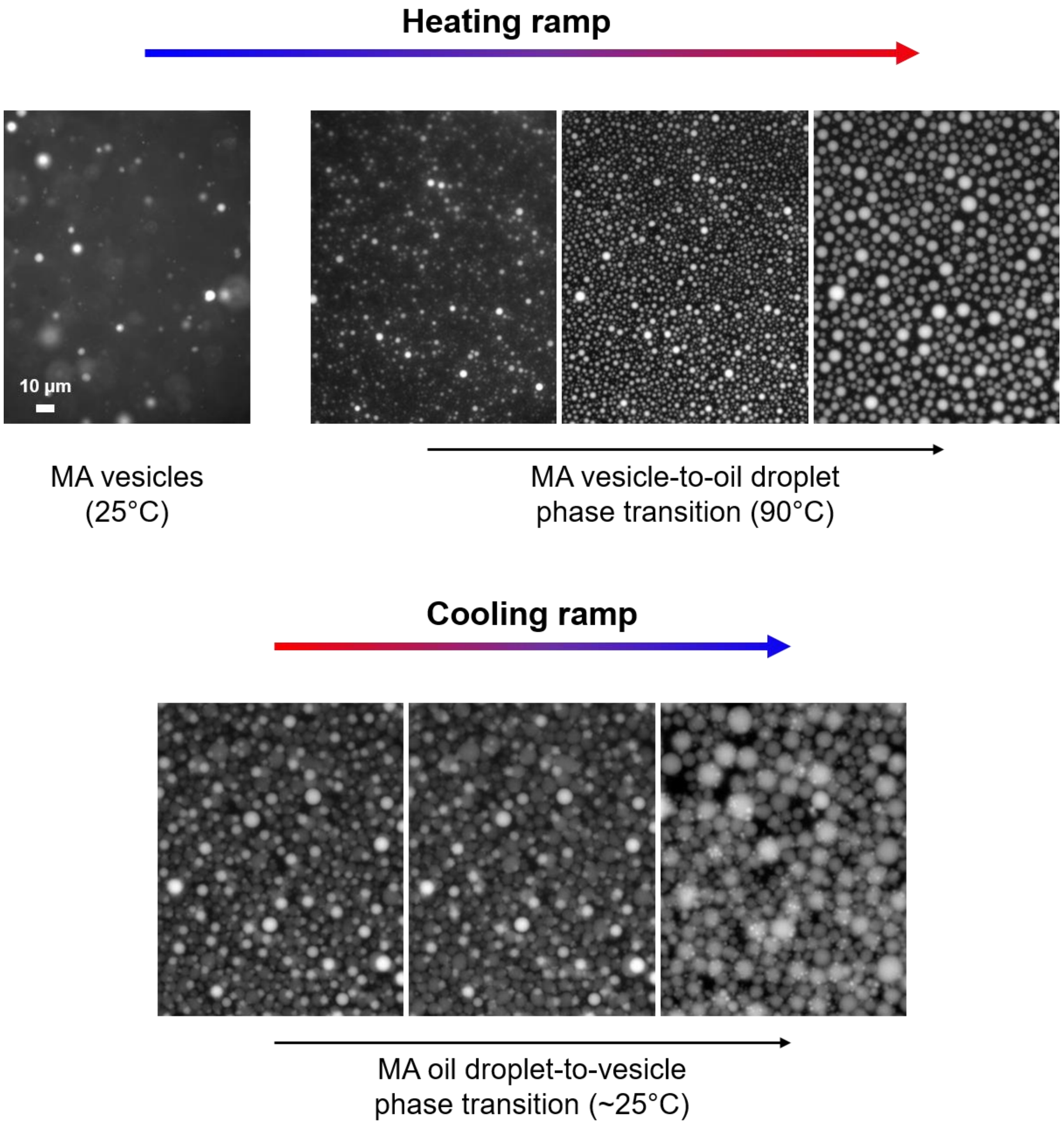


*Figure S14 – Frames extracted from movie S2, recorded during hot stage epifluorescence microscopy, for myristoleic acid vesicles. Once the phase transition temperature is reached (~80°C), myristoleic acid vesicles (10 mM) collapse into small oil droplets, which merge while kept at high temperature. The size of these oil droplets is smaller compared to the size of oil droplets observed for 50 mM myristoleic acid vesicles. During the cooling ramp, membrane budding can be observed from the surface of oil droplets (top). The size of these vesicles is also smaller compared to the size of re-generated vesicles observed for 50 mM myristoleic acid vesicles (bottom).*

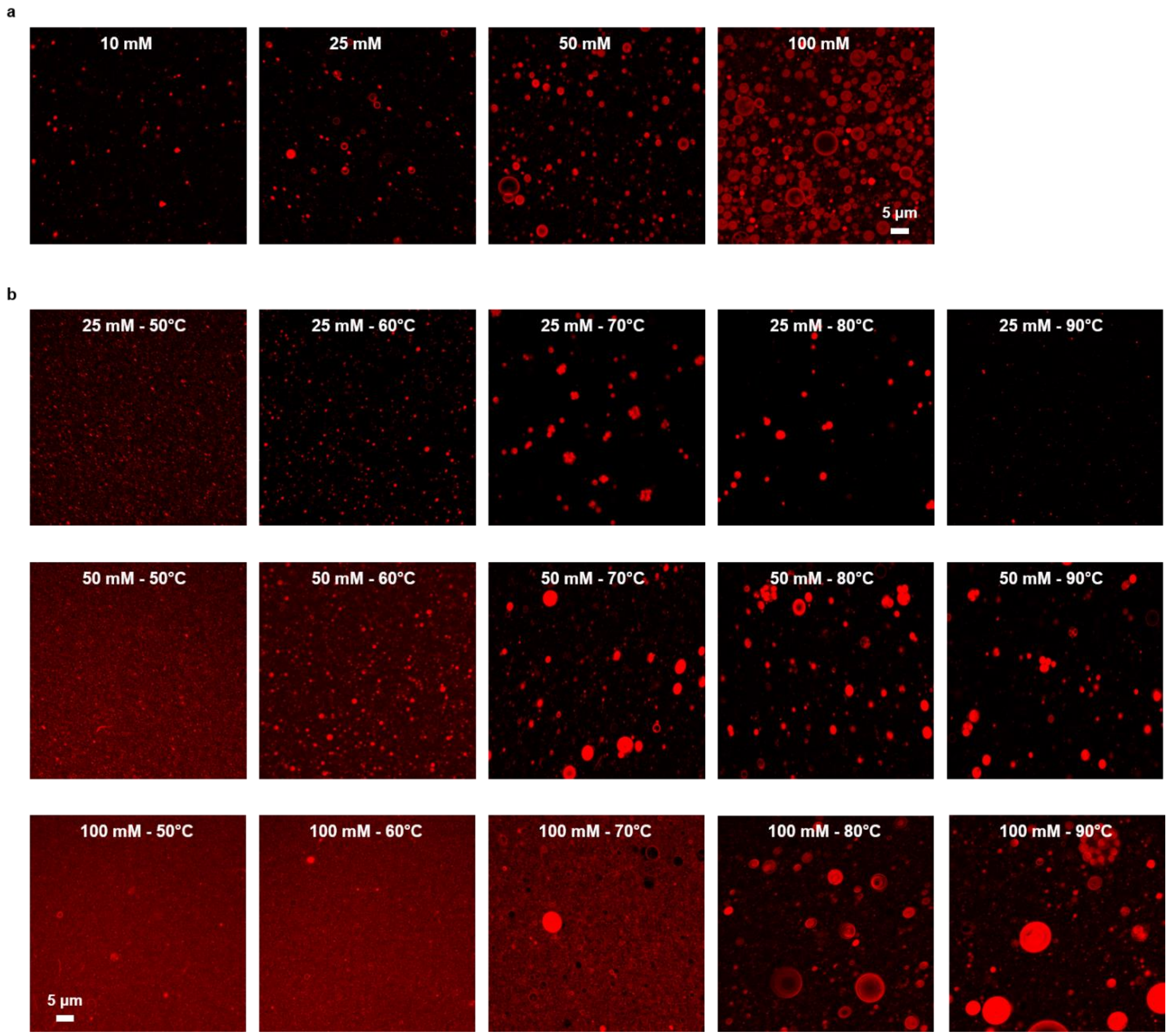


*Figure S15 – Representative confocal microscopy images used for statistical analysis. a) Confocal microscopy images taken for 100 nm-radius vesicles made of myristoleic acid (100 mM, 50 mM, 25 mM or 10 mM) in 200 mM Tris-HCl, pH 8 after heating them up to 90°C, and then cooling them down to 25°C and re-equilibrating them overnight. The increase in number of particles and size is observable. b) Confocal microscopy images taken for 100 nm-radius vesicles made of decanoic acid:decanol (2:1 ratio, 100 mM, 50 mM or 25 mM) in 200 mM Tris-HCl, pH 8 after heating them up to different temperatures (50°C, 60°C, 70°C, 80°C and 90°C) , and then cooling them down to 25°C and re-equilibrating them overnight. A similar trend to that monitored through absorbance profiles can be observed, confirming that phase transition occurs at higher temperatures for higher fatty acid concentrations. n = 10 replicates.*

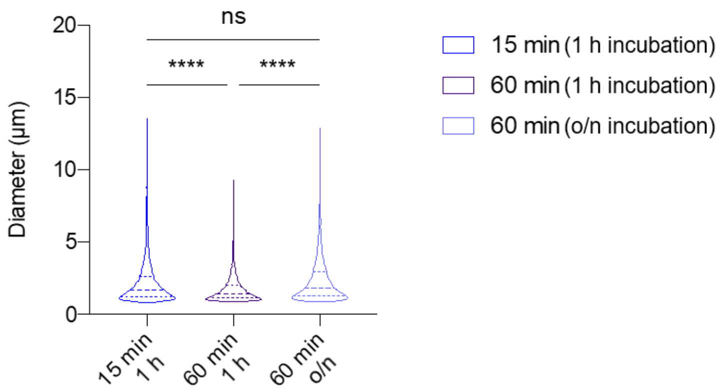


*Figure S16 – Violin plots of myristoleic acid samples showing how the size of next-generation vesicles is affected by re-equilibration time. Analyses were performed on collected confocal microscopy images for 100 nm-radius vesicles made of 50 mM myristoleic acid in 200 mM Tris-HCl, pH 8 after thermal cycling. Vesicles were heated up to 90°C and kept at high temperature for either 15 min or 60 min. After cooling the samples down to 25°C, vesicles were allowed to re-equilibrate for either 1 h or overnight. No significant differences could be observed for samples first heated for 15 min, and then cooled and re-equilibrated for either 1 h or overnight (data not shown). Statistical significance was assessed using the one-way ANOVA test, n = 10 replicates. Statistical values: ns, not significant; **** P = <0.0001.*

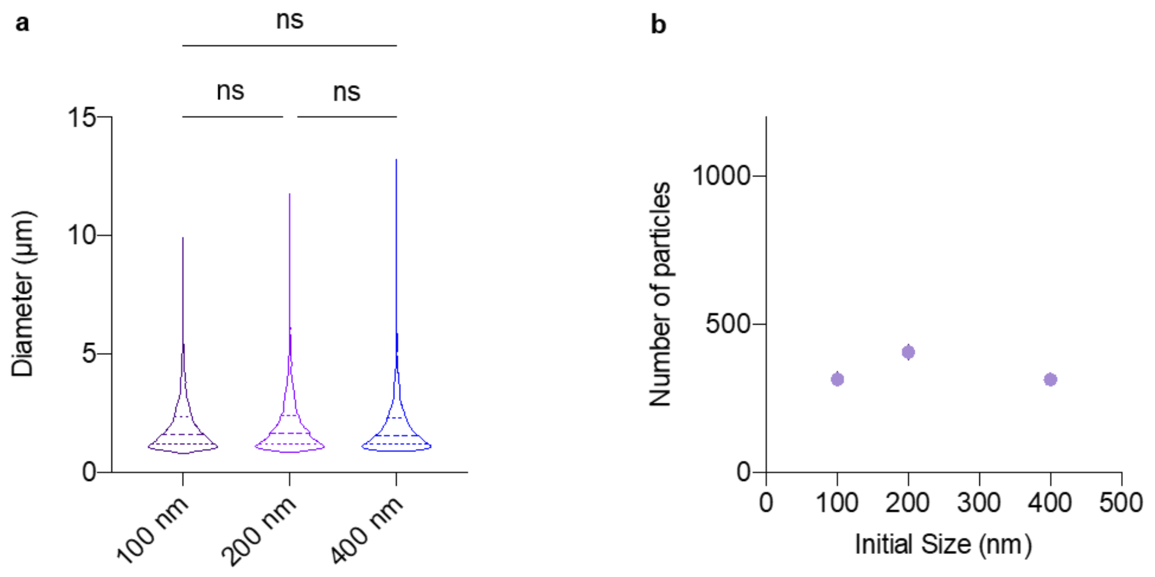


*Figure S17 – Analyses of myristoleic acid samples showing how the size and number of regenerated vesicles are affected by initial vesicle size. a) Violin plots showing the size distribution of regenerated vesicles. b) Vesicles number vs initial size graph for myristoleic acid vesicles. Analyses were performed on collected confocal microscopy images for vesicles (100 nm-, 200 nm- or 400 nm-diameter) made of 50 mM myristoleic acid in 200 mM Tris-HCl, pH 8 after thermal cycling. Vesicles were heated up to 90°C and kept at high temperature for 15 min. After cooling the samples down to 25°C, vesicles were allowed to re-equilibrate for 1 h. No significative differences could be observed when the samples were statistically compared. Statistical significance was assessed using the one-way ANOVA test, n = 10 replicates. Statistical values: ns, not significant.*

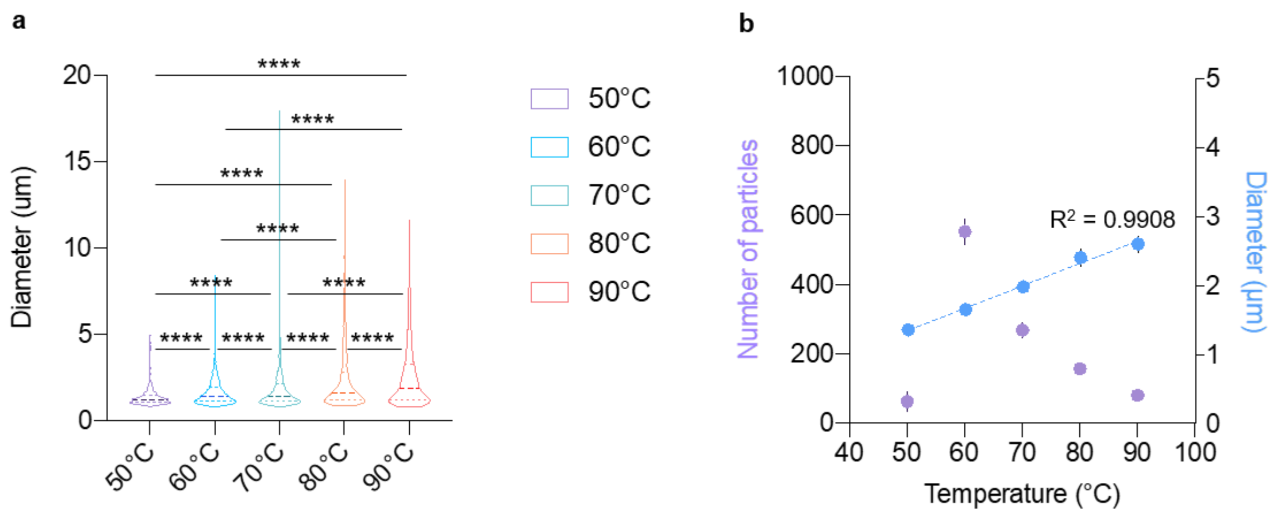


*Figure S18 – Analyses of decanoic acid-based samples showing how the size and number of regenerated vesicles are affected by maximum heating temperature. a) Violin plots showing the size distribution of regenerated vesicles. b) Vesicles number vs temperature and vesicle size vs temperature graphs for decanoic acid-based vesicles. Analyses were performed on collected confocal microscopy images for 100 nm-radius vesicles made of 50 mM decanoic acid:decanol (2:1 ratio) in 200 mM Tris-HCl, pH 8 after thermal cycling. Vesicles were heated up to 50°C, 60°C, 70°C, 80°C or 90°C and kept at high temperature for 15 min. After cooling the samples down to 25°C, vesicles were allowed to re-equilibrate for 1 h. Statistical significance was assessed using the one-way ANOVA test, n = 10 replicates. Statistical values for: **** P = <0.0001.*

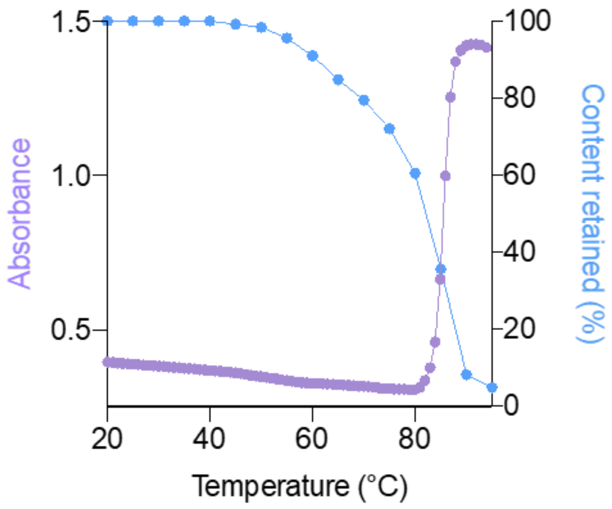


*Figure S19 – Overlapped profiles of turbidity and content retention as a function of temperature for 100 nm-radius vesicles made of 50 mM myristoleic acid vesicles containing 1 mM FITC-dextran, in 200 mM Tris-HCl, pH 8. Absorbance (light violet) is monitored at 420 nm, whereas content retention (light blue) for monitored by fluorescence ($\lambda_{exc}$ = 495 nm). The increase in turbidity perfectly matches a decrease in content retention. n = 3 replicates.*

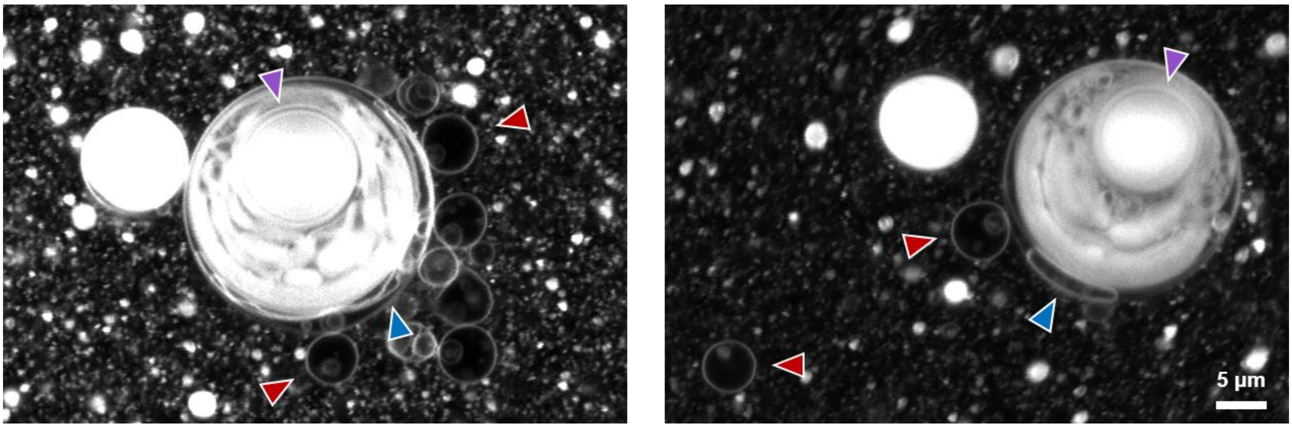


*Figure S20 – Representative confocal microscopy images showing the process of membrane formation from oil droplets, when samples are not allowed to re-equilibrate for enough time. Samples are representative for 100 nm-radius vesicles made of 50 mM myristoleic acid in 200 mM Tris-HCl, pH 8, after heating them up to 90°C for 1 h, and then cooling them down to 25°C and re-equilibrating them for 1 h. Violet arrows indicate sub-compartments of oil droplets within giant oil droplets. Blue arrows indicate fatty acid vesicles during the formation process from the external fatty acid layers of giant oil droplets. Red arrows indicate newly generated fatty acid vesicles.*

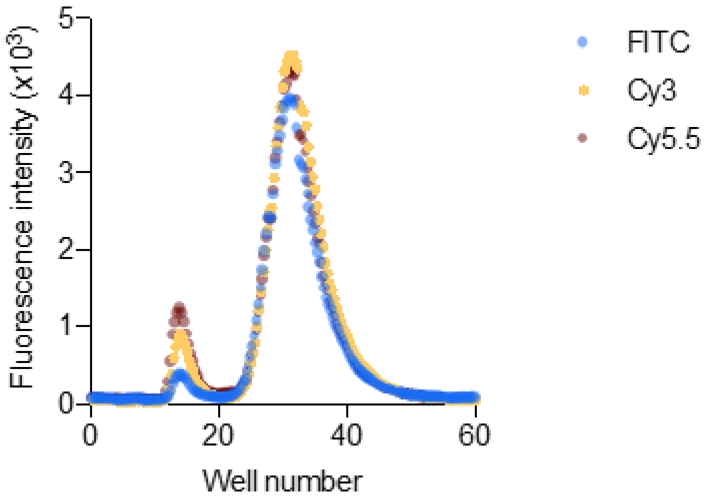


*Figure S21 – Size-exclusion chromatograms showing partial encapsulation of fluorescently-labelled 10-nt oligonucleotides in 100 nm-radius vesicles made of 50 mM myristoleic acid in 200 mM Tris-HCl, pH 8, upon heat exposure. Content uptake was monitored by fluorescence for FITC ($\lambda_{exc}$ = 495 nm), Cy3 ($\lambda_{exc}$ = 555 nm) and Cy5.5 ($\lambda_{exc}$ = 678 nm). n = 3 replicates.*

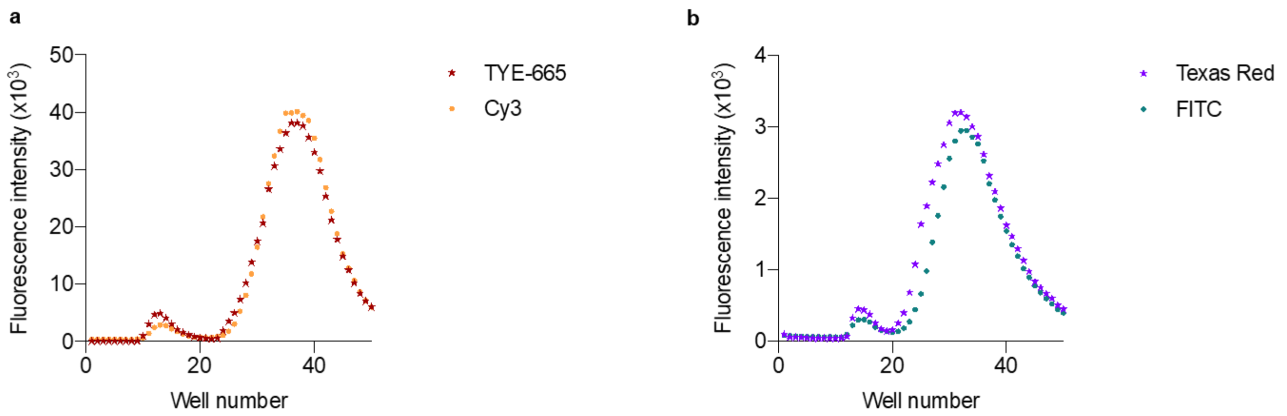


*Figure S22 – Size-exclusion chromatograms showing partial re-encapsulation of fluorescently-labelled extran or 10-nt oligonucleotides in 100 nm-radius vesicles made of 50 mM myristoleic acid in 200 mM Tris-HCl, pH 8, upon heat exposure. Content uptake was monitored by fluorescence for a) Cy3-10nt oligonucleotide ($\lambda_{exc}$ = 555 nm) and TYE665-10nt oligonucleotide ($\lambda_{exc}$ = 645 nm) and for b) FITC-dextran ($\lambda_{exc}$ = 495 nm) and Texas Red-dextran ($\lambda_{exc}$ = 580 nm). n = 3 replicates.*

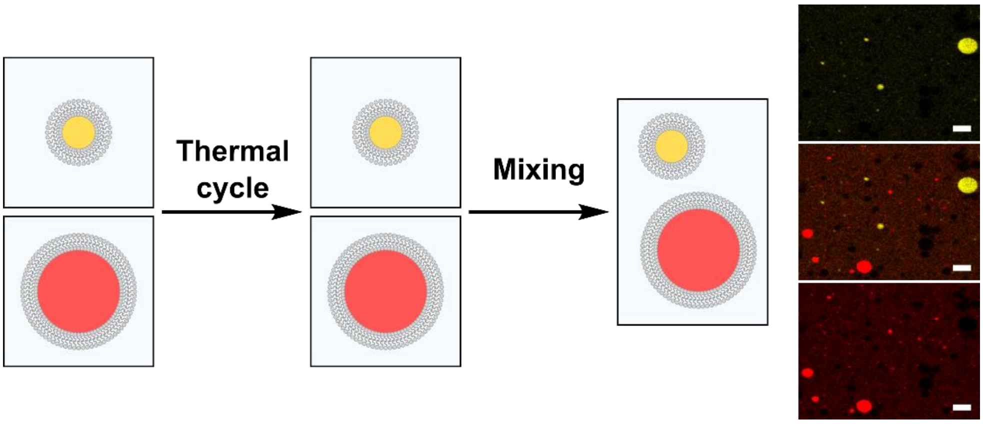


*Figure S23 – Schematic representation and confocal microscopy images for experiments with populations of vesicles with different fluorescent content. 100 nm-radius vesicles, made of 50 mM myristoleic acid in 200 mM Tris-HCl, pH 8 and containing either FITC-10nt or TYE665-10nt oligonucleotides, underwent thermal cycling and, after 1 h re-equilibration at room temperature, were mixed. Confocal microscopy images show no content mixing (top: FITC channel, bottom: TYE665 channel, middle: merged). Scale bar corresponds to 5 μm.*

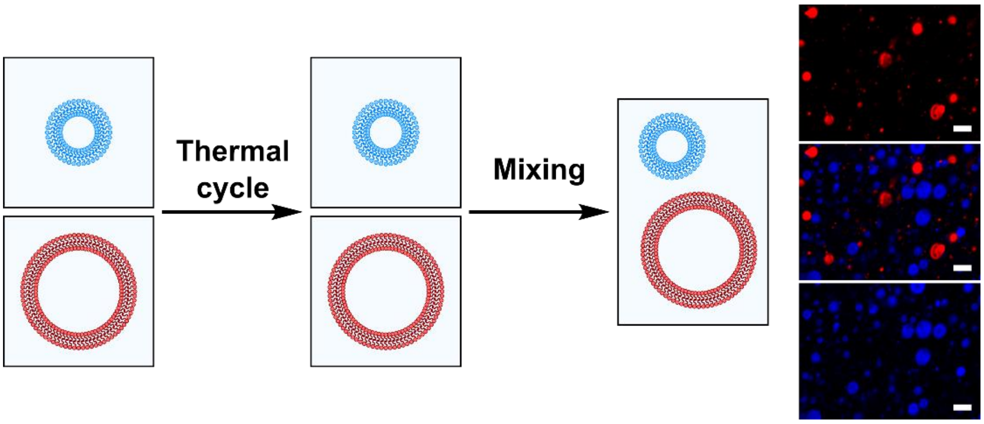


*Figure S24 – Schematic representation and confocal microscopy images for experiments with populations of vesicles, labelled with different fluorescent lipids. 100 nm-radius vesicles, made of 50 mM myristoleic acid in 200 mM Tris-HCl, pH 8 and either NBD-PE or Rh-DHPE, underwent thermal cycling and, after 1 h re-equilibration at room temperature, were mixed. Confocal microscopy images show no lipid mixing (top: Rh-DHPE channel, bottom: NBD-PE channel, middle: merged). Scale bar corresponds to 5 µm.*
